# Annotation of glycolysis, gluconeogenesis, and trehaloneogenesis pathways provide insight into carbohydrate metabolism in the Asian citrus psyllid

**DOI:** 10.1101/2021.10.11.463922

**Authors:** Blessy Tamayo, Kyle Kercher, Chad Vosburg, Crissy Massimino, Margaryta R. Jernigan, Denisse L. Hasan, Douglas Harper, Anuja Mathew, Samuel Adkins, Teresa Shippy, Prashant S. Hosmani, Mirella Flores-Gonzalez, Naftali Panitz, Lukas A. Mueller, Wayne B. Hunter, Joshua B. Benoit, Susan J. Brown, Tom D’Elia, Surya Saha

## Abstract

Citrus greening disease is caused by the pathogen *Candidatus* Liberibacter asiaticus, which is transmitted by the Asian citrus psyllid, *Diaphorina citri*. There is no curative treatment or significant prevention mechanism for this detrimental disease that causes continued economic losses from reduced citrus production. A high quality genome of *D. citri* is being manually annotated to provide accurate gene models required to identify novel control targets and increase understanding of this pest. Here, we annotated genes involved in glycolysis, gluconeogenesis, and trehaloneogenesis in the *D. citri* genome, as these are core metabolic pathways and suppression could reduce this pest. Specifically, twenty-five genes were identified and annotated in the glycolysis and gluconeogenesis pathways and seven genes for the trehaloneogenesis pathway. Comparative analysis showed that the glycolysis genes in *D. citri* are highly conserved compared to orthologs in other insect systems, but copy numbers vary in *D. citri*.

Expression levels of the annotated gene models were analyzed and several enzymes in the glycolysis pathway showed high expression in the thorax. This is consistent with the primary use of glucose by flight muscles located in the thorax. A few of the genes annotated in *D. citri* have been targeted for gene knockdown as a proof of concept, for RNAi therapeutics. Thus, manual annotation of these core metabolic pathways provides accurate genomic foundations for developing gene-targeting therapeutics to reduce *D. citri*.

## Introduction

Huanglongbing, HLB, or citrus greening disease is the biggest global threat to the citrus industry throughout the world [1]. The phloem-limited bacterial pathogen *Candidatus* Liberibacter asiaticus (*C*Las) is the causative agent of HLB. Upon infection of a citrus tree, HLB causes development of small, bitter fruits, loss of tree vigor, fruit drop, and ultimately tree decline and death [1]; [2]; [3], [4]. This bacterium is transmitted by the psyllid vector, *D. citri*, when feeding on citrus [5]; [6]. Pesticide application to eliminate *D. citri* has not been successful and no cure for HLB exists [7]; [8]. To develop new psyllid control strategies, The International Psyllid Genome Consortium was established in 2009 [9] to provide the genome, transcriptome resources, and an Official Gene Set of *D. citri* [10]; [11]. A recent, nearly complete genome with significantly improved gene accuracy has been generated, providing a significant data set for the establishment of gene-targeted strategies to suppress psyllid populations (*opensource*: Diaci_v3.0, www.citrusgreening.org) [USDA-NIFA grant 2015-70016-23028]. As part of this genome project, we conducted manual annotation of genes in critical pathways to provide the quality gene models required for designing molecular therapeutics such as RNA interference (RNAi) [12]; [13]; [14]; [15]; [16]; [17]; [18]; [19]; [20], antisense oligonucleotides (ASO) [21]; [15]; [19], and gene editing (CRISPR) [22]; [23]. Here, we examined *D. citri* orthologs associated with the critical metabolic pathways glycolysis, gluconeogenesis, and trehaloneogenesis.

## Context

A community-driven annotation strategy was used to identify and characterize the genes encoding enzymes involved in glycolysis, gluconeogenesis, and trehaloneogenesis (Fig. 1). Glycolysis is a major metabolic pathway that plays a vital role in core energy processing reactions, and as a source of metabolites for other biochemical processes. For insects, there is a high utilization of glucose in flight muscle located in the thorax [24]. As a result, the activities of glycolytic enzymes are increased in insect flight muscle compared to vertebrate muscle tissue [25]. Gluconeogenesis is essential in insects to maintain sugar homeostasis and serves as the initial step towards the generation of glucose disaccharide, also known as trehalose. Trehalose is the main circulating sugar in the insect hemolymph [26]; [27]; [28]. In trehaloneogenesis, glucose- 6-phosphate is converted into trehalose by trehalose 6 phosphate synthase (TPS).

**Figure 1:**
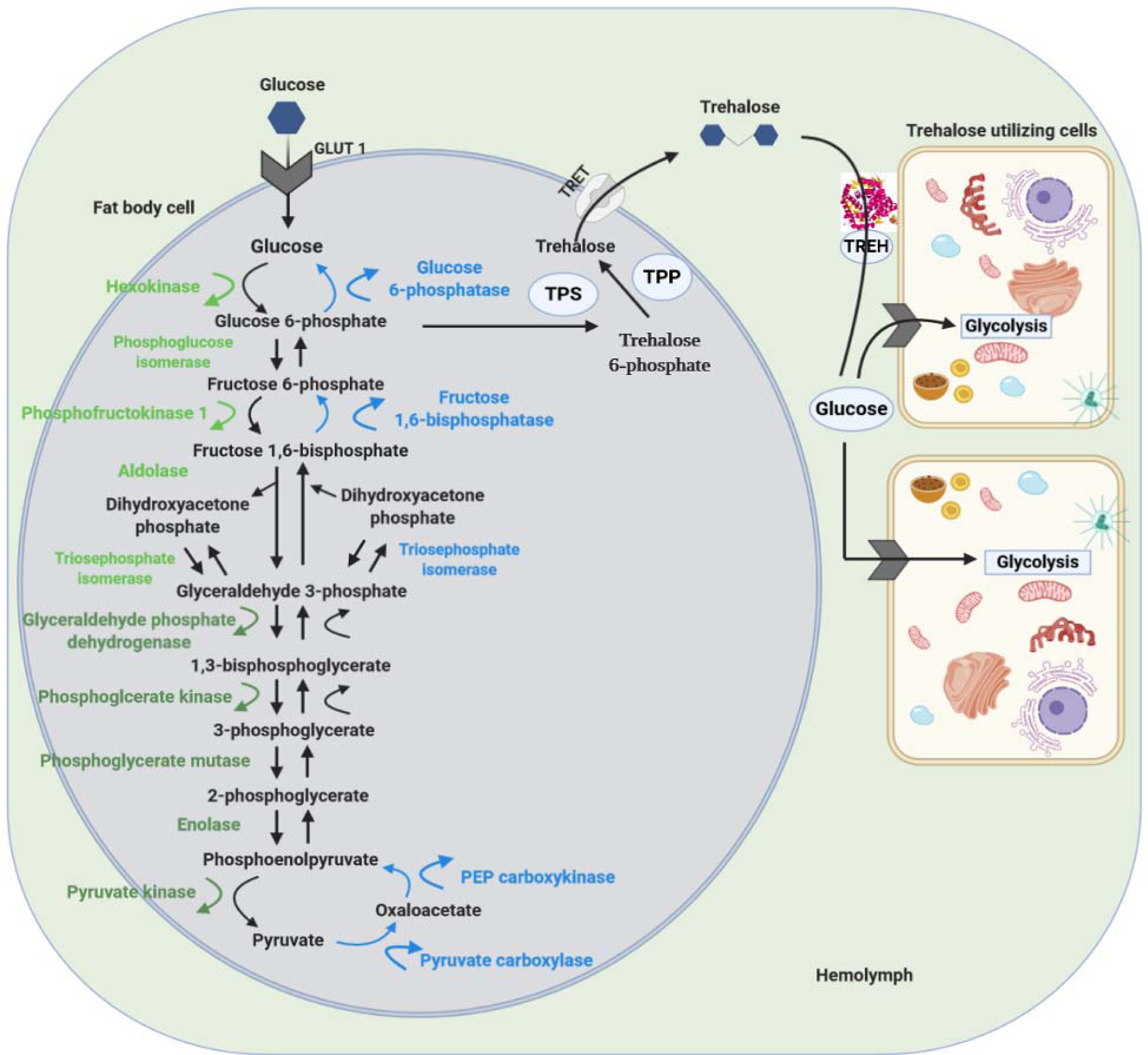
Overview of Glycolysis, Gluconeogenesis, and Trehaloneogenesis Pathway Image.

Trehalase enzymes then degrade trehalose into two glucose molecules [29]. Genes involved in psyllid glycolysis, gluconeogenesis, and trehaloneogenesis have been targeted by several RNAi studies (Appendix Table 1) as a promising avenue for psyllid population suppression. In particular, one proof of concept experiment targeting trehalase has led to the release of the first RNAi patent to control psyllid populations [30]. RNAi, as a biopesticide, and strategies for delivery and applications to target insect pests and viral pathogens have been thoroughly reviewed [31]; [32]; [33]; [34]; [35].

The pathway image shows the enzymes that produce and utilize glucose and trehalose in insects [36]. The glycolysis pathway consists of ten enzymes that convert glucose into pyruvate as the final product. These are divided into the energy investment phase (light green) and the energy production phase (dark green). The gluconeogenesis pathway consists of eight enzymes (blue) with three being unique to the pathway that bypasses the irreversible reactions in glycolysis to convert non-carbohydrate molecules into glucose. The trehaloneogenesis pathway consists of three enzymes which are *Trehalose-6-phosphate synthase* (*TPS*), *Trehalose-6-phosphate phosphatase* (*TPP*), and *Trehalase* (*TREH*), as well as *Trehalose transporters* (*TRET*) and *Glucose transporters* (*GLUT 1*). Image was adapted from a diagram in [37], and was created with BioRender.com.

## Methods

The *D. citri* genome was manually annotated through a collaborative community [11] driven strategy with an undergraduate focus that allows specific students to focus on main gene sets [38]. Orthologous protein sequences for the glycolysis, gluconeogenesis, and trehaloneogenesis pathways were obtained from the National Center for Biotechnology Information (NCBI) protein database [39] and were used to BLAST the *D. citri* MCOT (Maker (RRID:SCR_005309), Cufflinks (RRID:SCR_014597), Oases (RRID:SCR_011896), and Trinity (RRID:SCR_013048)) protein database to find predicted protein models [36]. The MCOT predicted protein models were used for searching the *D. citri* genomes (version 2.0 and 3.0) [38]. Regions of high sequence identity were manually curated in Apollo version 2.1.0 (RRID:SCR_001936) using *de novo* transcriptome, MCOT gene predictions, RNA-seq, Iso-seq, and ortholog data to support and evaluate gene structure (Appendix Table 2). The curated gene models were compared to other orthologous sequences, such as hemipterans, available through NCBI for accuracy. A more detailed description of the annotation workflow is available (Fig. 2) [40].

**Figure 2:**
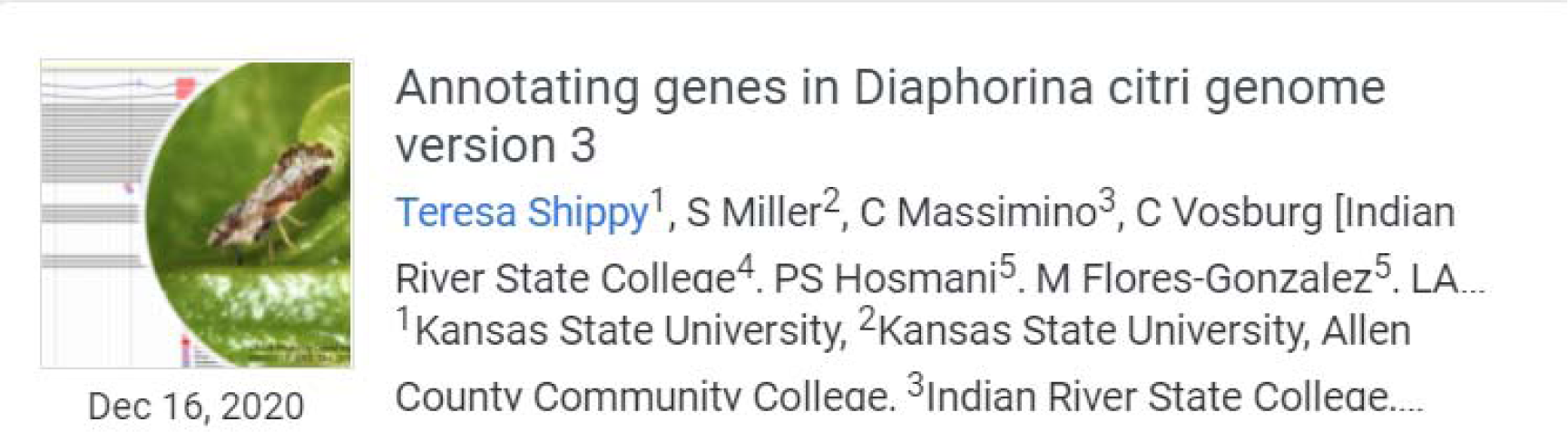
Protocols.io protocol for psyllid genome curation [40].

Neighbor-joining phylogenetic tree of the annotated *hexokinase* gene models in *D. citri* and orthologous sequences were created with MEGA version 7 (RRID:SCR_000667) using the MUSCLE (RRID:SCR_011812) multiple sequence alignment with p-distance for determining branch length and 1,000 bootstrap replicates [41].

Expression levels of the carbohydrate metabolism genes throughout different life stages (egg, nymph, and adult) in *C*Las infected and uninfected *D. citri* insects were collected from the Citrus Greening Expression Network (CGEN) [36] and visualized using Excel (RRID:SCR_016137) and the pheatmap package in R. (RRID:SCR:_016418).

### Data Validation and Quality Control

The characterization of the carbohydrate metabolism pathways in *D. citri* is separated into four sections: the energy investment phase of glycolysis, the energy production phase of glycolysis, gluconeogenesis, and trehaloneogenesis. The enzymes involved in the breakdown and synthesis of glucose and trehalose were annotated in version 3.0 of the *D. citri* genome [42]. The following genes in the energy investment phase: *hexokinase* (*HK*), *phosphoglucose isomerase* (*PGI*), *phosphofructokinase*(*PFK*), *Fructose-bisphosphate aldolase* (*aldolase*), *triosephosphate isomerase* (*TPI*), and in the energy production phase: *glyceraldehyde phosphate dehydrogenase* (*GAPDH*), *phosphoglycerate kinase* (*PGK*), *phosphoglycerate mutase* (*PGAM*), *enolase*, and *pyruvate kinase* (*PYK*) were annotated. The annotated genes for gluconeogenesis are *pyruvate carboxylase* (*PC*), *phosphoenolpyruvate carboxykinase* (*PEPCK*), and *fructose 1,6-bisphosphatase* (*FBPase*). In trehaloneogenesis, *trehalose transporter 1* (*TRET-1*) and *2* (*TRET-2*), *glucose transporter 1* (*GLUT-1*), and two gene models of both *trehalose-6-phosphate synthase* (*TPS*) and *trehalase* (*TREH*) were annotated. Gene expression data sets in CGEN were analyzed for potential differences, as expression patterns can provide insight into potential RNAi target candidates for molecular therapeutics (Appendix Table 1).

Orthologous sequences from related insects and information about conserved motifs or domains were used to determine the final annotation. We used proteins from *Drosophila melanogaster* (*Dm*) [43], *Tribolium castaneum* (*Tc*) [44], *Apis mellifera*(*Am*) [45], *Acyrthosiphon pisum* (*Ap*) [46], *Nilaparvata lugens* (*Nl*) [47]; [48], *Halyomorpha halys* (*Hh*) [49].

### Energy investment phase of glycolysis

*HK* catalyzes the first step in glycolysis, utilizing ATP to phosphorylate glucose creating glucose-6-phosphate. Most insects have multiple *HK* genes and three copies of *HK* are present in the *D. citri* genome (Fig. 3, Table 3 and Appendix Table 2). In insect flight muscles, *HK* activity is inhibited by its product, glucose-6-phosphate, to initiate flight muscle activity [50]. *Drosophila melanogaster* has four duplicated *HK* genes, with *Hex-A* being the most conserved and essential flight muscle *HK* isozyme among *Drosophila* species [51], [52]. For *D. citri*, one of the copies of *HK* type 2-2 (Dcitr03g19430.1.1) showed moderate expression in the male and female thorax. In contrast, another copy *HK* type 2-3 (Dcitr06g14200.1.1), was highly expressed in the adult gut and midgut when compared to *HK* type 2-2 and its overall expression (Fig. 4)

**Figure 3:**
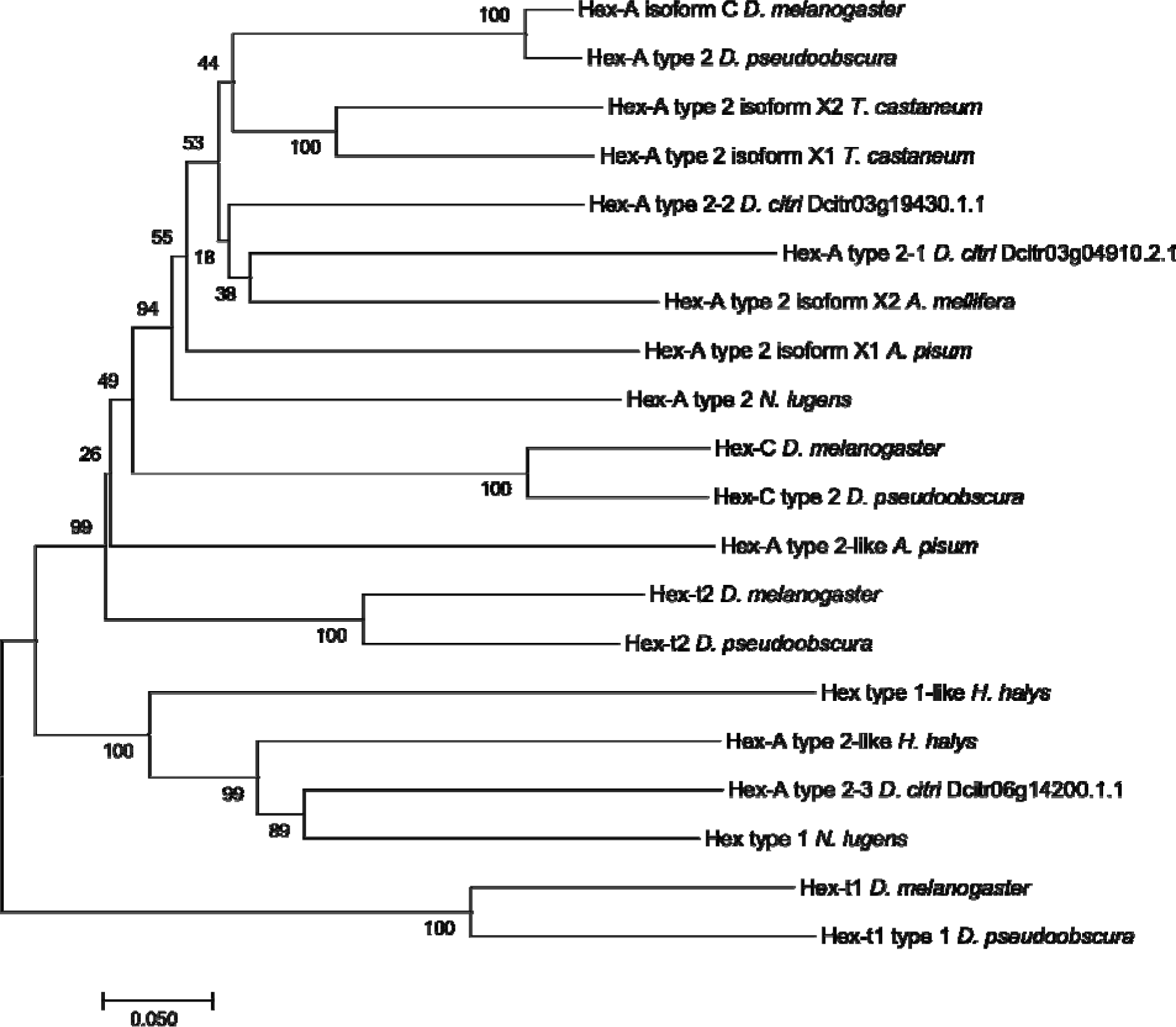
Phylogenetic analysis of *hexokinase* (*HK*). *Hexokinase* amino acid sequence of *D. citri* compared with sequences from other insects. MUSCLE multiple sequence alignments of *HK* in *D. citri* and orthologs were performed on MEGA7 and neighbor joining phylogenetic trees were constructed with p-distance for determining evolutionary distance and 1000 bootstrapping replicates [41]. The accession numbers of the orthologous sequences used in the phylogenetic analysis can be located in Table 1.

**Table 1:** Accession Numbers.

| Protein | *Drosophila melanogaster* | *Tribolium castaneum* | *Apis mellifera* | *Acyrtosiphon pisum* | *Drosophila pseudoobscura* | *Halyomorpha halys* | *Nilaparvata lugens* |
| --- | --- | --- | --- | --- | --- | --- | --- |
| Hex-A | NP_001259<br>384.1 | XP_0082<br>01714.1<br>XP_9706<br>45.1 | XP_006<br>557646.<br>1 | XP_0032422<br>38.1<br>XP_0019524<br>12.1 | XP_0013550<br>83.1 |  | XP_022204<br>875.1 |
| Hex-type 1 |  |  |  |  |  | XP_0142822<br>49.1 | XP_022184<br>109.1 |
| Hex-type 2 |  |  |  |  |  | XP_0142827<br>21.1 |  |
| Hex-C | NP_524674.<br>1 |  |  |  | XP_0013601<br>04.2 |  |  |
| Hex-t1 | NP_788744.<br>1 |  |  |  | XP_0013591<br>46.2 |  |  |
| Hex-t2 | NP_733151.<br>2 |  |  |  | XP_0021376<br>41.2 |  |  |

The accession numbers for *hexokinase* orthologs were obtained from the NCBI database.

**Figure 4:**
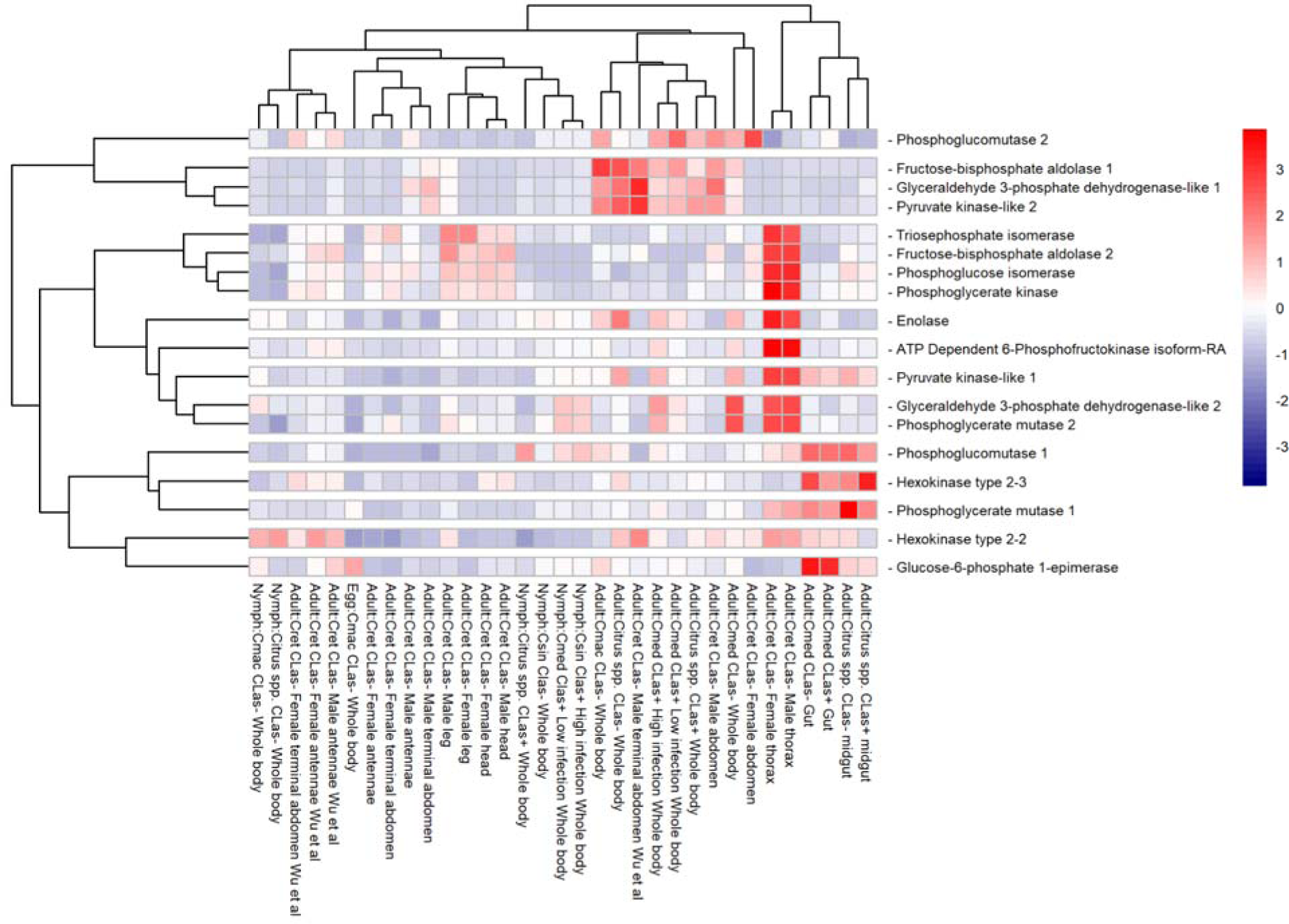
Comparison of RNA-Seq datasets of genes involved in glycolysis. The heatmap shows results from *D. citri* reared on various citrus varieties, both infected and uninfected with *C*Las. Expression values were collected from CGEN [36]. Data in the heatmap shows transcripts per million scaled by gene. RNA-seq data is available from NCBI Bioproject’s PRJNA609978 and PRJNA448935 and in addition to several published data sets [9], [53] [54], [55], [56]. Expression data for *HK type 2-1* (Dcitr03g04910.2.1) is not present in the heatmap.

**Table 3:**
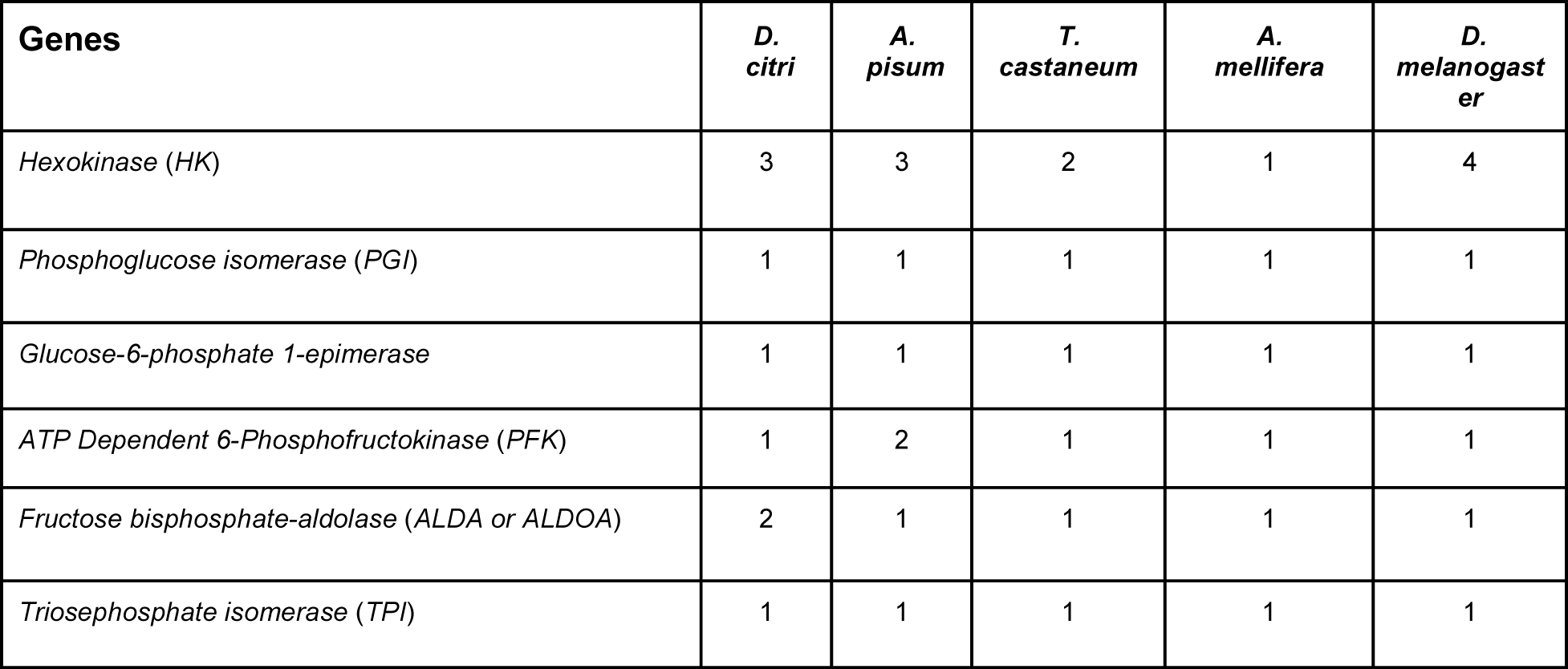
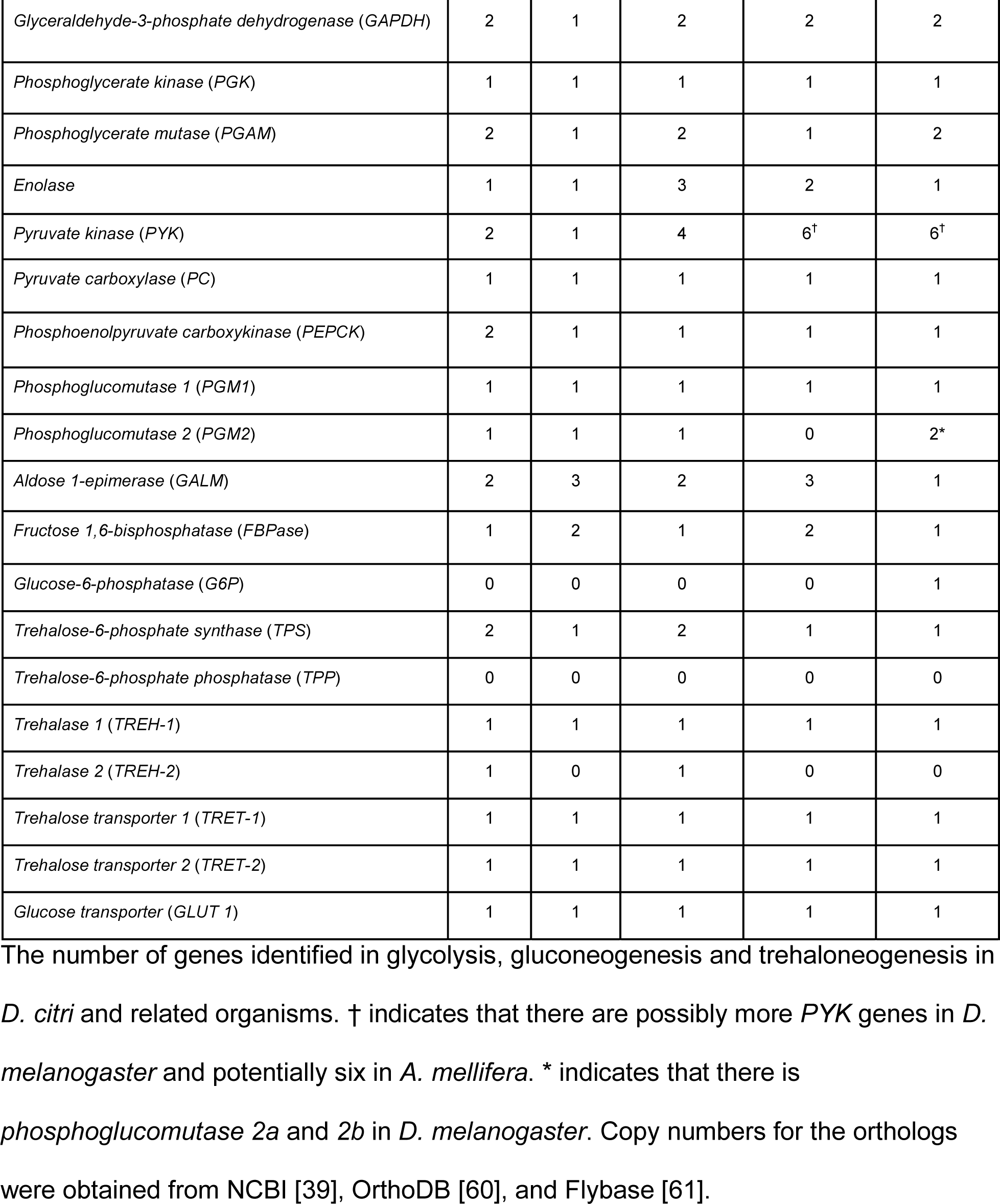
Gene Copy Number.

| Genes | *D. citri* | *A. pisum* | *T. castaneum* | *A. mellifera* | *D. melanogaster* |
| --- | --- | --- | --- | --- | --- |
| *Hexokinase (HK)* | 3 | 3 | 2 | 1 | 4 |
| *Phosphoglucose isomerase (PGI)* | 1 | 1 | 1 | 1 | 1 |
| *Glucose-6-phosphate 1-epimerase* | 1 | 1 | 1 | 1 | 1 |
| *ATP Dependent 6-Phosphofructokinase (PFK)* | 1 | 2 | 1 | 1 | 1 |
| *Fructose bisphosphate-aldolase (ALDA or ALDOA)* | 2 | 1 | 1 | 1 | 1 |
| *Triosephosphate isomerase (TPI)* | 1 | 1 | 1 | 1 | 1 |
| *Glyceraldehyde-3-phosphate dehydrogenase (GAPDH)* | 2 | 1 | 2 | 2 | 2 |
| *Phosphoglycerate kinase (PGK)* | 1 | 1 | 1 | 1 | 1 |
| *Phosphoglycerate mutase (PGAM)* | 2 | 1 | 2 | 1 | 2 |
| *Enolase* | 1 | 1 | 3 | 2 | 1 |
| *Pyruvate kinase (PYK)* | 2 | 1 | 4 | 6<sup>†</sup> | 6<sup>†</sup> |
| *Pyruvate carboxylase (PC)* | 1 | 1 | 1 | 1 | 1 |
| *Phosphoenolpyruvate carboxykinase (PEPCK)* | 2 | 1 | 1 | 1 | 1 |
| *Phosphoglucomutase 1 (PGM1)* | 1 | 1 | 1 | 1 | 1 |
| *Phosphoglucomutase 2 (PGM2)* | 1 | 1 | 1 | 0 | 2* |
| *Aldose 1-epimerase (GALM)* | 2 | 3 | 2 | 3 | 1 |
| *Fructose 1,6-bisphosphatase (FBPase)* | 1 | 2 | 1 | 2 | 1 |
| *Glucose-6-phosphatase (G6P)* | 0 | 0 | 0 | 0 | 1 |
| *Trehalose-6-phosphate synthase (TPS)* | 2 | 1 | 2 | 1 | 1 |
| *Trehalose-6-phosphate phosphatase (TPP)* | 0 | 0 | 0 | 0 | 0 |
| *Trehalase 1 (TREH-1)* | 1 | 1 | 1 | 1 | 1 |
| *Trehalase 2 (TREH-2)* | 1 | 0 | 1 | 0 | 0 |
| *Trehalose transporter 1 (TRET-1)* | 1 | 1 | 1 | 1 | 1 |
| *Trehalose transporter 2 (TRET-2)* | 1 | 1 | 1 | 1 | 1 |
| *Glucose transporter (GLUT 1)* | 1 | 1 | 1 | 1 | 1 |

The number of genes identified in glycolysis, gluconeogenesis and trehaloneogenesis in *D. citri* and related organisms. † indicates that there are possibly more *PYK* genes in *D. melanogaster* and potentially six in *A. mellifera*. \* indicates that there is *phosphoglucomutase 2a* and *2b* in *D. melanogaster*. Copy numbers for the orthologs were obtained from NCBI [39], OrthoDB [60], and Flybase [61].

*PGI* catalyzes the interconversion of glucose-6-phosphate and fructose-6-phosphate in the second step of glycolysis. Consistent with the gene copy number of *PGI* for orthologs in other insects, such as *D. melanogaster, Apis mellifera*, *Acyrthosiphon pisum*, and *Tribolium castaneum,* a single copy of *PGI* (Dcitr00g06460.1.1) was found. Expression for *PGI* is high in the male and female thorax (Fig. 4).

*PFK*, which catalyzes the phosphorylation of fructose-6-phosphate using ATP to generate fructose-1,6-bisphosphate and ADP, is the key regulatory enzyme controlling glycolysis in insects, as it catalyzes a rate-determining reaction [57], [58]. One copy of *PFK* (Dcitr01g16570.1.1) was found and annotated in *D. citri* (Table 3). *Aldolase* catalyzes the fourth step, the reversible aldol cleavage of fructose-1,6-bisphosphate to form two trioses, glyceraldehyde-3-phosphate (GAP) and dihydroxyacetone phosphate (DHAP). Although most insects have a single copy of this gene, two well supported copies were found in *D. citri* (Table 3 and 4). One of the *aldolase* annotated copies, *fructose-bisphosphate aldolase 1*, (Dcitr04g02510.1.1) appears to have moderate expression in the male abdomen and terminal abdomen and highest expression in the adult whole body (Fig. 4). *TPI* catalyzes the fifth step, the reversible interconversion of DHAP and GAP. *TPI* is also important to sustain DHAP to maintain insect flight muscle activity [59]. *D. citri* contains a single copy of this gene (Dcitr10g08030.1.1), which is consistent with other insects (Table 3). Expression of several of these genes in the investment phase were shown to be high in the male and female thorax, especially in *PFK* (Dcitr01g16570.1.1), *fructose-bisphosphate aldolase 2* (Dcitr11g09140.1.1), and *TPI* (Dcitr10g08030.1.1) (NCBI BioProject PRJNA448935) (Fig. 4, Appendix Table 3).

### Energy production phase of Glycolysis

*GAPDH* catalyzes the reversible conversion of GAP to 1,3-bisphosphoglycerate during glycolysis. Two *GAPDH* genes were annotated in *D. citri* and the expression data for the two paralogs show that *GAPDH-like 1* (Dcitr10g11030.1.1) has higher expression in the male terminal abdomen and whole body and *GAPDH-like 2* (Dcitr01g03200.1.1) has higher expression values overall with a considerable increase in male thorax, female thorax and whole body (NCBI BioProject PRJNA609978, NCBI BioProject PRJNA448935) (Fig. 4, Appendix Table 4). *PGK* catalyzes the reversible conversion of 1,3-bisphosphoglycerate to 3-phosphoglycerate (3PG) while generating one molecule of ATP in the seventh step of glycolysis. A single gene was annotated in *D. citri*, and other insects also have single copies (Table 3). *PGAM* is an enzyme that converts 3- phosphoglycerate to 2-phosphoglycerate. Members of the *PGAM* family share a common *PGAM* domain, and function as either phosphotransferases or phosphohydrolases [62]. Two copies of *PGAM* were annotated in the *D. citri* genome (Table 3). *PGAM 1* (Dcitr03g11640.1.1) has high expression evident in the midgut and the other paralog, *PGAM 2* (Dcitr03g17850.1.1) is highly expressed in the whole body (Fig. 4).

*Enolase* catalyzes the conversion of 2-phosphoglycerate to phosphoenolpyruvate in the ninth step of the glycolytic pathway and a single copy was annotated in the *D. citri* genome (Table 3). RNAi knockdown of the *-enolase* in *Nilaparvata lugens* reduced egg α production, offspring, and hatching rate; however, mortality of adults was unaffected [63]. Pairwise alignment between the *N. lugens* and *D. citri* sequences reveal the characteristics of the *enolase* family: a hydrophobic domain (AAVPSGASTGI) in the N-terminal region at position 31-41, a seven amino acid substrate binding pocket (H159, E211, K345, HRS373-375, and K396), a metal-binding site (S38, D246, E295, and D320) and the *enolase* signature motif (LLLKVNQIGSVTES) [63].

*PYK* catalyzes the irreversible transfer of a phosphoryl group from phosphoenolpyruvate to ADP; thus generating pyruvate and a second ATP molecule, the end products of the glycolysis reaction. The copy number of *PYK* varies among insects; *A. mellifera* and *D. melanogaster* both contain six, and *A. gambiae* has only one (Table 3). In *D. citri,* two *PYK* genes were characterized and annotated (Appendix Table 2). One of the *PYK* genes (Dcitr07g06140.1.1) is highly expressed in male and female thorax and the other *PYK* gene (Dcitr01g11190.1.1) has relatively low overall expression with the highest expression in the male terminal abdomen (Fig. 4).

Expression analysis of the enzymes from this phase of glycolysis in thoracic tissue shows that the highest expression is observed for *GAPDH-like 2* and *PYK-like 1* and the lowest occurs for both *GAPDH-like 1* and *PYK-like 2* (Fig. 5). In addition, *PGK* (Dcitr00g01740.1.1) and *enolase* (Dcitr02g07600.1.1) also have high expression in the male and female thorax and *PGAM2* (Dcitr03g17850.1.1) has high expression in whole body besides the male and female thorax (NCBI BioProject PRJNA609978, NCBI BioProject PRJNA448935) (Fig. 4, Appendix Table 4).

**Figure 5:**
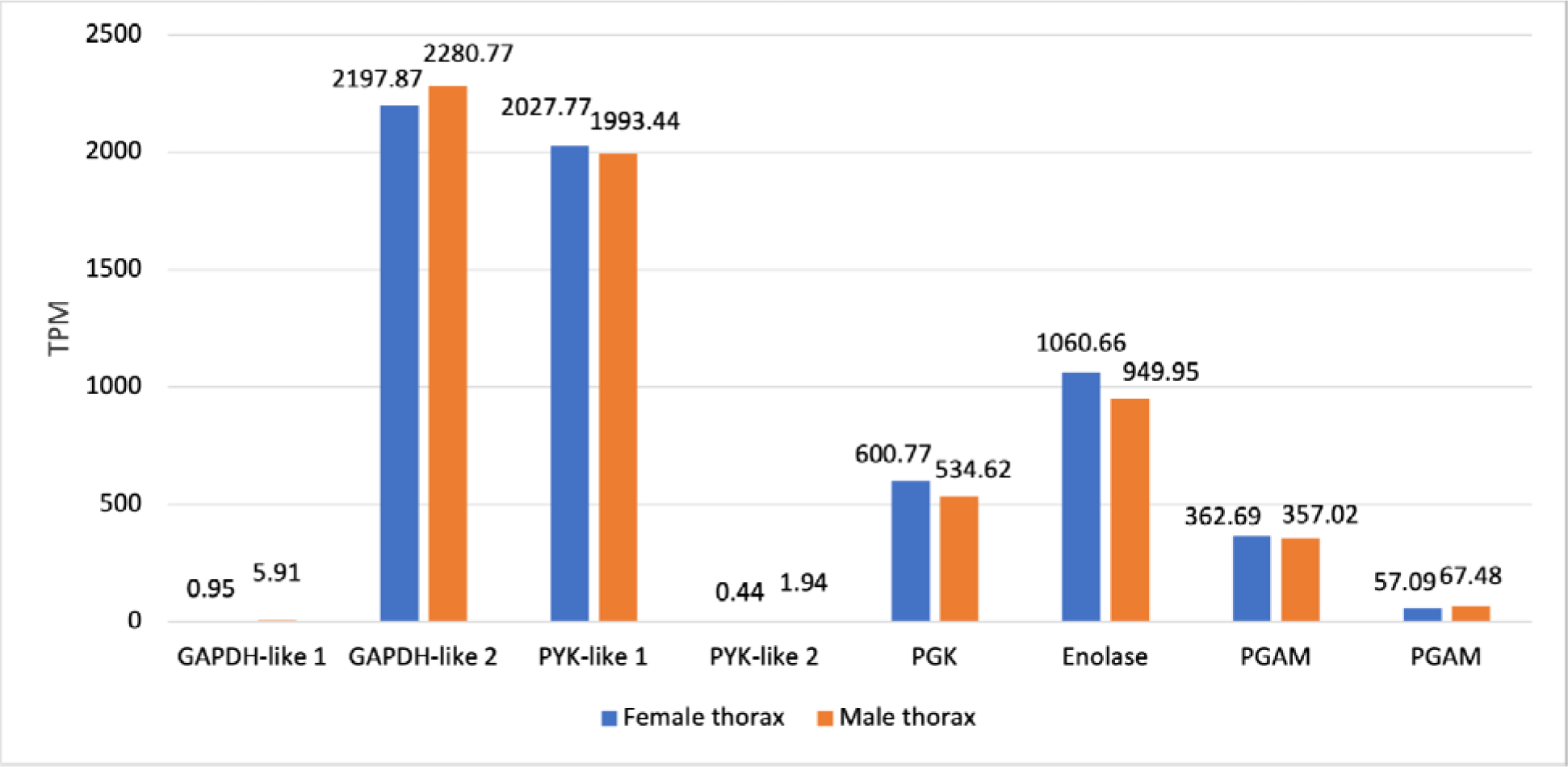
PEN expression data for the enzymes involved during the energy production phase in *D. citri.* (*GAPDH-like 1*: Dcitr10g11030.1.1; *GAPDH-like 2*: Dcitr01g03200.1.1; *PYK-like 1*: Dcitr07g06140.1.1; *PYK-like 2*: Dcitr01g11190.1.1; *PGK*: Dcitr00g01740.1.1; *Enolase*: Dcitr02g07600.1.1; *PGAM*: Dcitr03g17850.1.1, Dcitr03g11640.1.1 respectively). Values are based on transcripts taken from the thorax of healthy *C*Las- *D. citri* male and female adults that fed on *C. reticulata*. These experiments had a single replicate. RNA-seq data is available from NCBI BioProject’s PRJNA448935.

### Enzymes of Gluconeogenesis

Gluconeogenesis is the metabolic process to re-generate glucose from non- carbohydrate substrates and uses four specific enzymes. *PC* catalyzes the ATP- dependent carboxylation of pyruvate to oxaloacetate. The curated *PC* model (Dcitr08g01610.1.1) in *D. citri* shows highest overall expression in the male and female thorax, male and female head, and male and female antenna (Fig. 6, Fig. 7, Appendix Table 5). *PEPCK* controls the cataplerotic flux and converts oxaloacetate from the tricarboxylic acid cycle to form PEP. Two *PEPCK* genes were annotated and characterized in the *D. citri* genome (Appendix Table 2). The first *PEPCK* copy (Dcitr05g10240.1.1) has the highest expression in most tissues compared to all of the gluconeogenesis genes as is evident in the male and female antenna, male and female thorax, and the male and female head. The highest expression of the second copy of *PEPCK* (Dcitr08g02760.1.1) is shown in the whole body. *FBPase* facilitates one of the three bypass reactions occurring in gluconeogenesis where the hydrolysis of fructose- 1,6-bisphosphate produces fructose-6-phosphate. A single copy of this gene was annotated in *D. citri,* which is comparable to other insects, although two copies are present in the pea aphid, *A. pisum,* and honey bee, *A. mellifera*(Table 3). *FBPase* (Dcitr11g08070.1.1) shows highest expression in the egg (Fig. 5). *Glucose-6- phosphatase* (*G6Pase* or *G6P*), which is specific to gluconeogenesis, catalyzes the conversion of glucose-6-phosphate to glucose [27]. However, this enzyme is not present in most insect species, including *D. citri.* Though present in *N. lugens*, RNAi studies showed that knockdown of *G6Pase* in *N. lugens* had no effect on the genes involved in trehalose metabolism [64].

**Figure 6:**
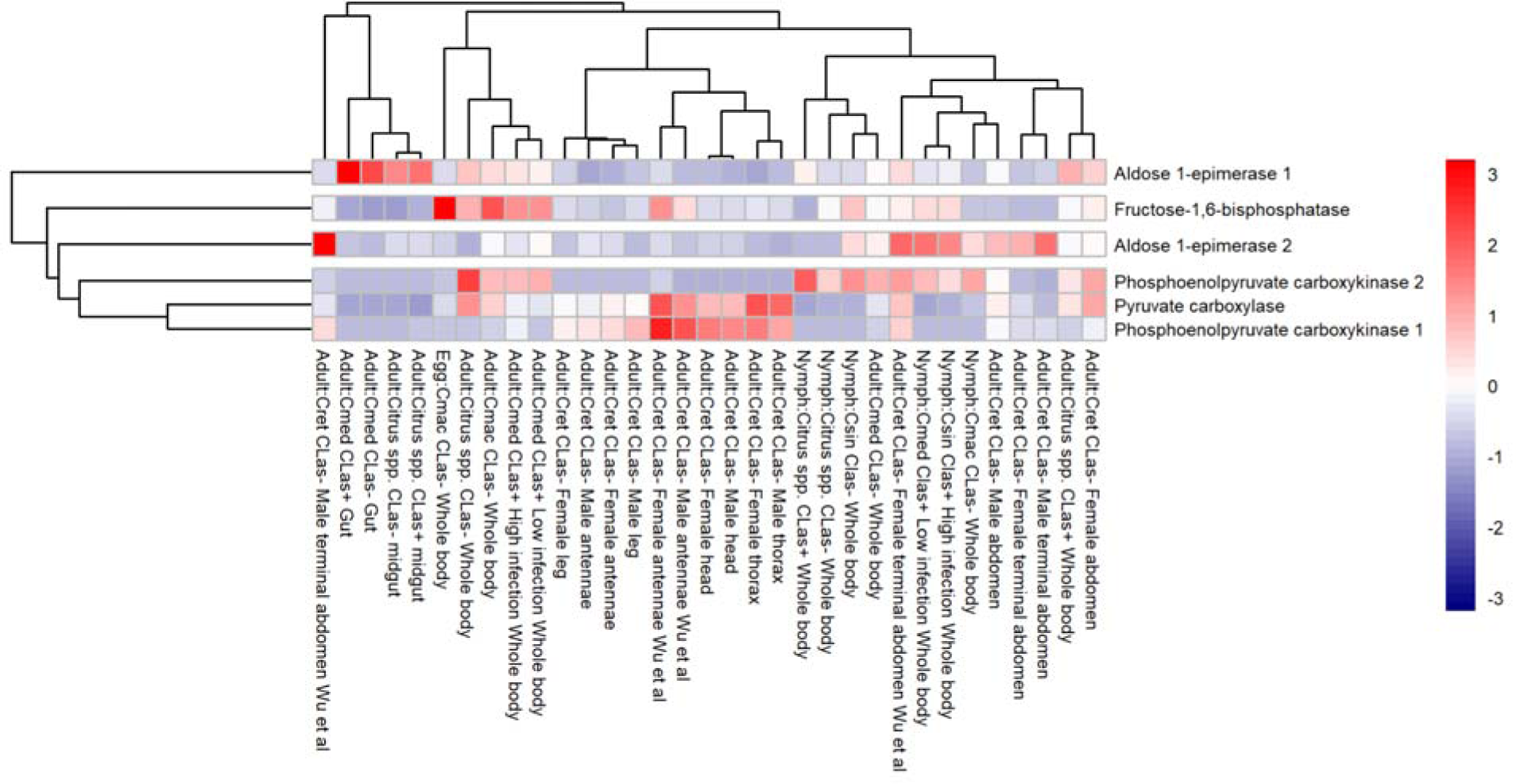
Comparison of RNA-Seq datasets of genes involved in gluconeogenesis. The heatmap shows results from *D. citri* reared on various citrus varieties, both infected and uninfected with *C*Las. Expression values were collected from Citrus Greening expression network [36]. Data in the heatmap shows transcripts per million scaled by gene. RNA-seq data is available from NCBI Bioprojects PRJNA609978 and PRJNA448935 and published data sets [53].

**Figure 7:**
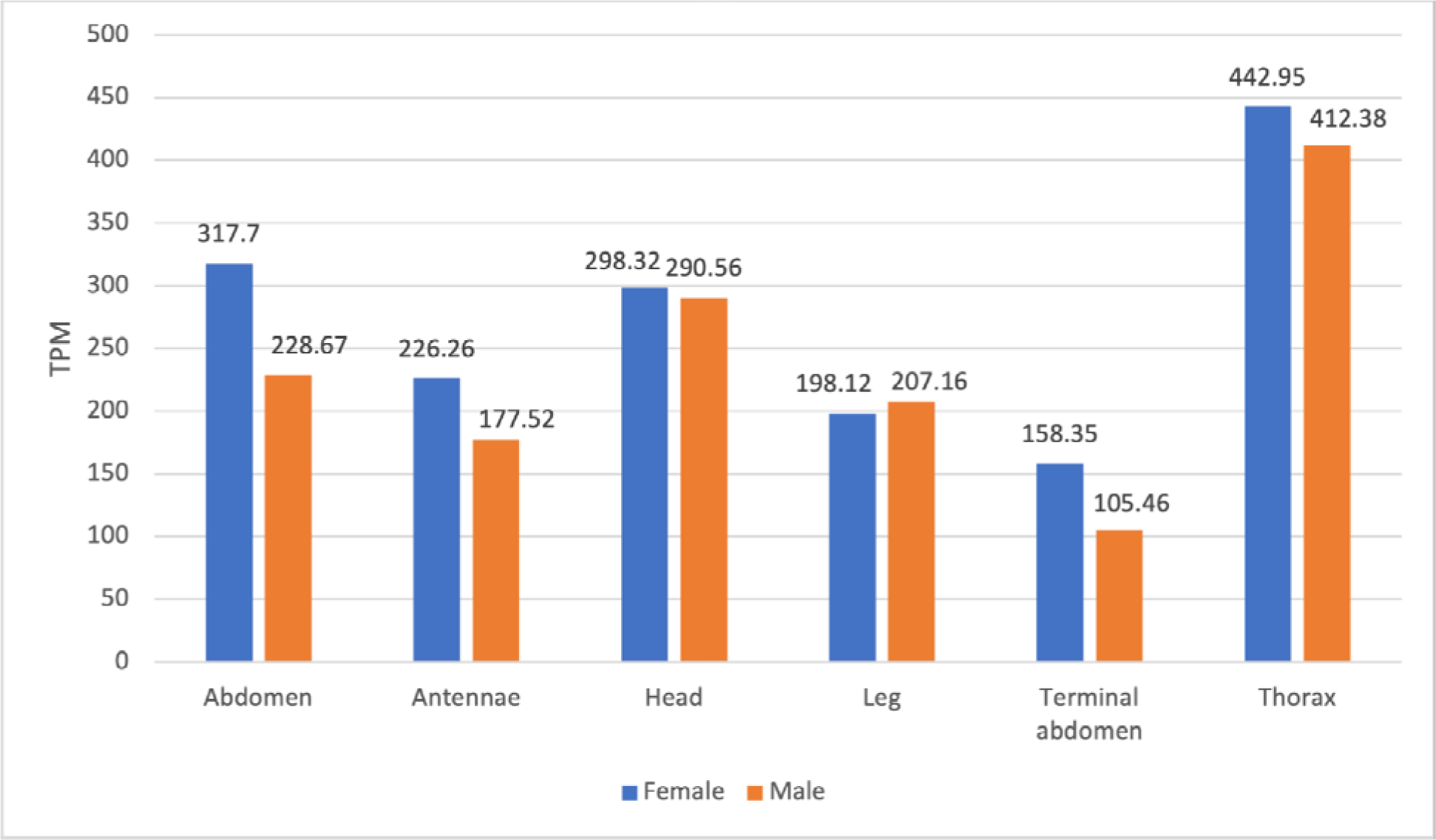
PEN Expression data of the enzyme *Pyruvate carboxylase* (Dcitr08g01610.1.1) in *D. citri*. Values are based on transcripts isolated from various body parts of healthy *C*Las- *D. citri* adults that fed on *C. reticulata.* These experiments had a single replicate. RNA-seq data is available from NCBI BioProjects PRJNA448935.

### Enzymes of Trehaloneogenesis

Trehalose is a nonreducing disaccharide that is present in many organisms, including yeast, fungi, bacteria, plants and invertebrates and is in high concentration as the main hemolymph sugar in insects [28], [65]. Trehalose is synthesized from glucose by trehalose-6-phosphate (Tre-6-P), where the mobilization of trehalose to glucose is considered critical for metabolic homeostasis in insect physiology [26]. Synthesis of trehalose occurs in the fat body, when stimulated by neuropeptides from the brain [28]. These peptides decrease the concentration of fructose 2,6-bisphosphate which strongly activates the glycolytic enzyme *PFK* and inhibits the gluconeogenic enzyme *fructose 1,6-bisphosphatase*. *Fructose 2,6-bisphosphatase* is thus a key metabolic signal in regulating trehalose synthesis in insects. After synthesis, trehalose is transported through the hemolymph and enters cells through *trehalose transporters*, where it is converted into glucose by trehalase.

There are three enzymes involved in trehaloneogenesis: *trehalose-6-phosphate synthase* (*TPS*), *trehalose-6-phosphate phosphatase* (*TPP*), and *trehalase* (*TREH*) (Fig. 1). *TPS* catalyzes the transfer of glucose from UDP-glucose to G6P forming trehalose 6-phosphate (T6P) and UDP [66]. A *TPS* (*DcTPS*) gene has been targeted in *D. citri* for RNAi therapeutics and the results suggested that dsRNA-mediated gene specific silencing resulted in a strong reduction in expression of *DcTPS* and survival rate of nymphs and an increase in malformation [67]. Two copies of *TPS* were annotated in the v3 genome of *D. citri*. *TPS 1* (Dcitr02g17550.1.1) had the highest expression found in the *C*Las+ and *C*Las- adult midgut, respectively (Fig. 9, Appendix Table 6). In some organisms, *TPP* dephosphorylates T6P to trehalose and inorganic phosphate [66].

However, many insects appear to lack this gene, including *D. citri* as it was not found in the v3 genome. Most insects with multiple *TPS* genes encode proteins with TPS and TPP domains [68], [69]. *TPS* in *Drosophila* appears to have the functions of both *TPS* and *TPP* [70]. *Trehalase* (*TREH*) catalyzes stored trehalose by cleaving it to two glucose molecules. There are two trehalase genes: *TREH-1*, which encodes a soluble enzyme found in haemolymph, goblet cell cavity and egg homogenates, and *TREH-2*, which encodes a membrane-bound enzyme found in flight muscle, ovary, spermatophore, midgut, brain and thoracic ganglia [66]. The two curated *TREH* genes in *D. citri* show different expression in the psyllid. *TREH-1A* (Dcitr07g04030.1.1) shows high expression in the gut and midgut and *TREH*-2 (Dcitr08g09220.1.1) shows moderate expression in the female thorax and male antennae (Fig. 8). *TREH* is the only enzyme known for the irreversible splitting of trehalose in all insects [66] and *D. citri and T. castaneum* are the only insects with the second copy, *TREH-2* (Table 3).

**Figure 8:**
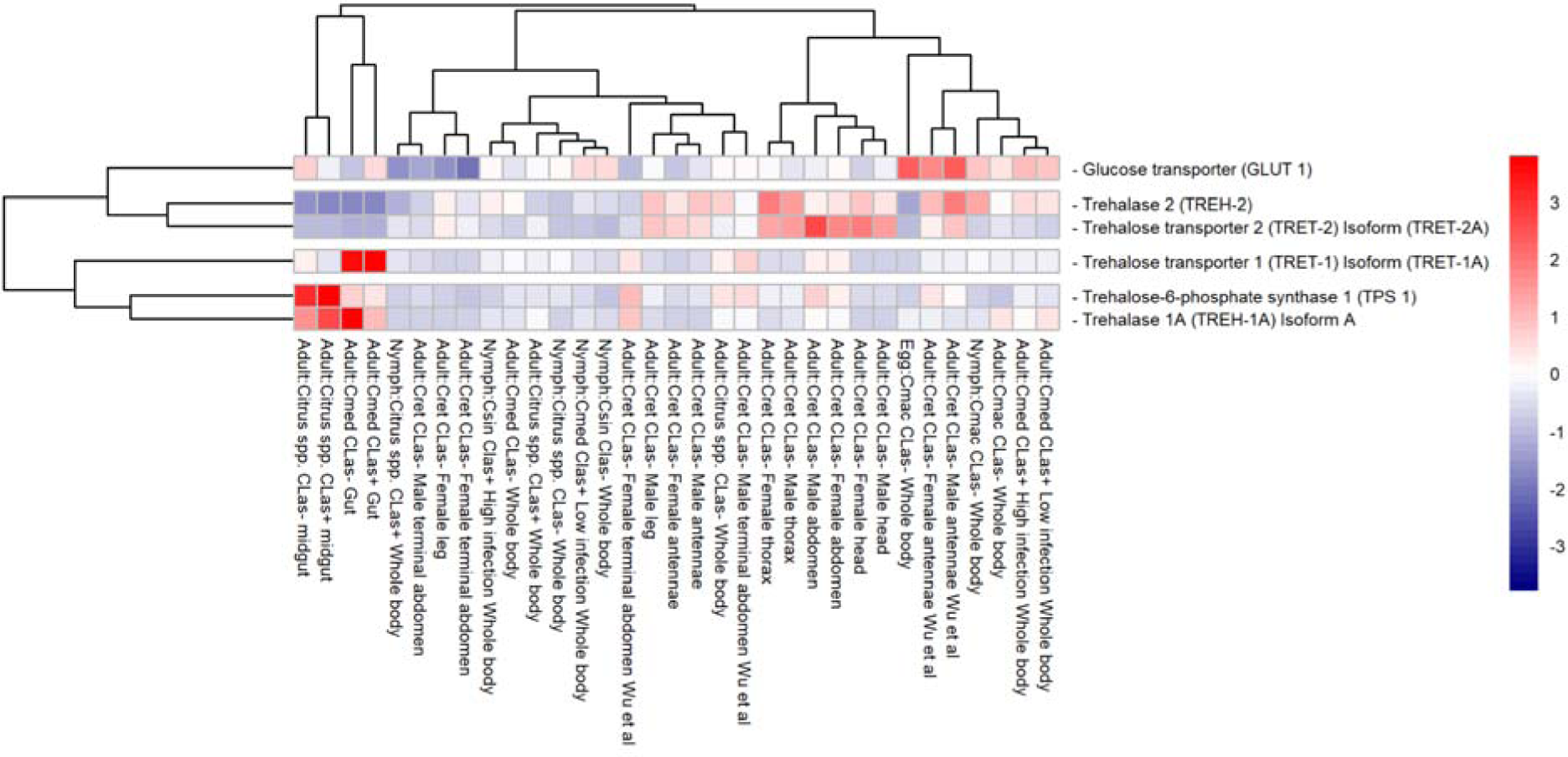
Comparison of RNA-Seq datasets of genes involved in trehaloneogenesis. The heatmap shows results from *D. citri* reared on various citrus varieties, both infected and uninfected with *C*Las. Expression values were collected from Citrus Greening expression network [36]. Data in the heatmap shows transcripts per million scaled by gene. RNA-seq data is available from NCBI Bioproject’s PRJNA609978 and PRJNA448935 and published data sets [53]. Expression data for *Trehalose-6- phosphate synthase 2* (*TPS 2*), *Trehalase 2* (*TREH-2*), *Trehalose transporter 1* (*TRET-1*) *Isoform* (*TRET-1B*), *Trehalose transporter 2* (*TRET-2*) *Isoform* (*TRET-2B*), and *Trehalose transporter 2* (*TRET-2*) *Isoform* (*TRET-2C*) are not present in the heatmap.

**Figure 9:**
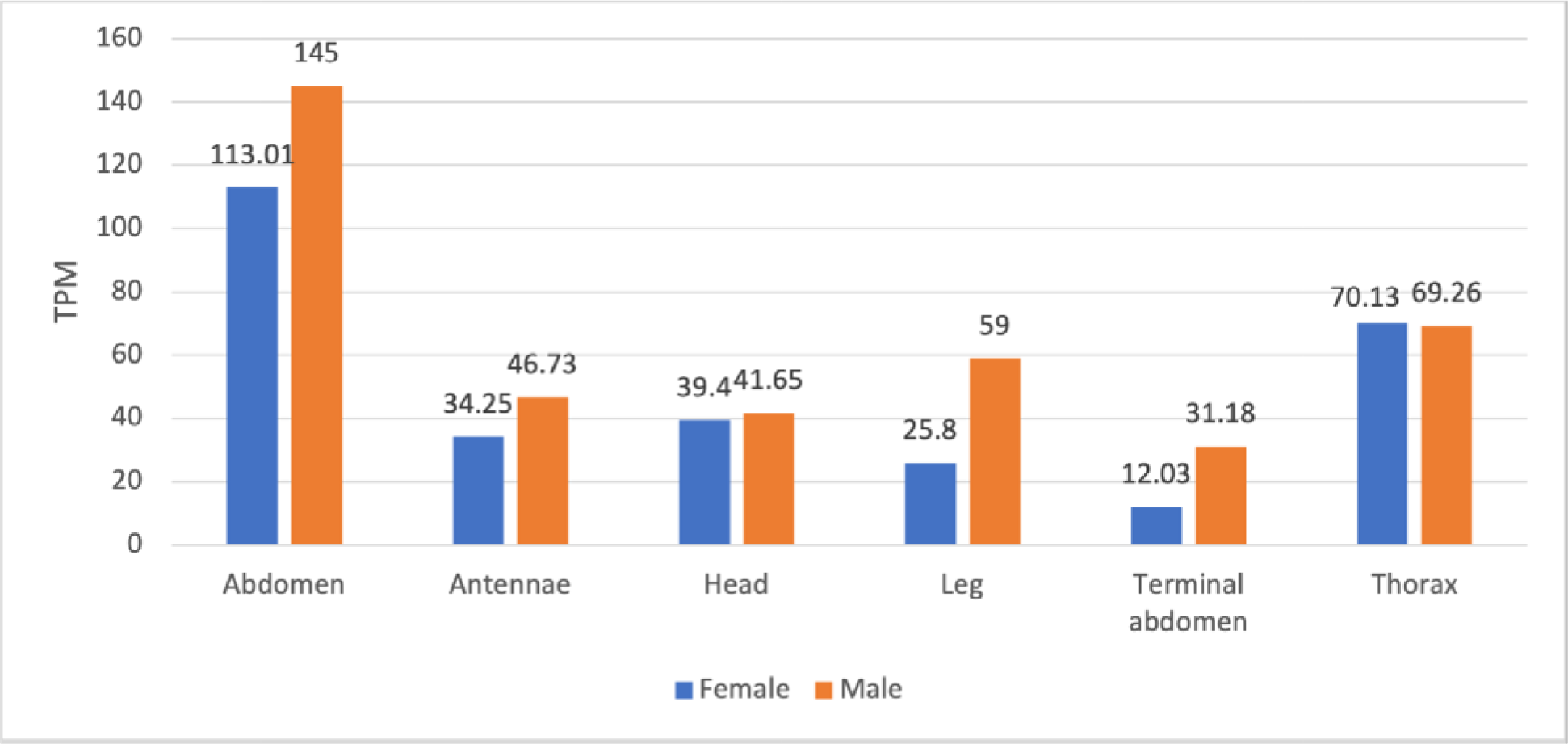
PEN Expression data of the enzyme *Trehalose 6-phosphate synthase* (Dcitr02g17550.1.1) in *D. citri*. Values are based on transcripts expressed in various body parts of healthy *C*Las- *D. citri* adults that fed on *C. reticulata*. These experiments had a single replicate. RNA-seq data is available from NCBI BioProject PRJNA448935.

The two main *trehalose transporters* are *trehalose transporter 1* (*TRET1*) and *trehalose transporter 2* (*TRET2*), which both transport trehalose to and from cells with *TREH*. One gene copy for each of these *trehalose transporters* was annotated in *D. citri* (Appendix Table 2) and expression analysis shows that *TRET1* (Dcitr01g17710.1.1) is highly expressed in the gut and *TRET2* (Dcitr00g03240.1.1) is moderately expressed in the male abdomen (Fig. 8).

## Conclusion

Manual curation of genes in the glycolysis, gluconeogenesis, and trehaloneogenesis pathways was completed in the genome of *D. citri*. Twenty-five genes were annotated in glycolysis and gluconeogenesis. The pathways are highly conserved and copy numbers of the genes annotated were comparable to other insects. Except for *G6Pase*, all enzymes involved in glycolysis and gluconeogenesis were identified. An additional seven genes involved in trehaloneogenesis were also identified and annotated. Manual annotation of these central metabolic pathways provides accurate gene models which are required for development of molecular therapeutics to target *D. citri*. RNAi studies targeting genes involved in trehalose metabolism produced significant mortality in *D. citri*, [67], [71] demonstrating the functional application of the genes identified. Expression analysis of the genes annotated in the carbohydrate metabolism pathways identified differences related to life stage, sex and tissue. This data advances the understanding of the basic biology of *D. citri* and will aid in the development of RNAi-based applications.

## Reuse potential

The manually curated gene models were annotated through a collaborative community project [11] to further understand psyllid biology and with a goal to annotate gene families related to immune response, metabolism and other major functions [72]. Continued examination of the glycolysis, gluconeogenesis, and trehaloneogenesis pathways across arthropods, and especially in insect vectors like *D. citri*, will provide novel and species-specific gene targets to control psyllid populations (potentially through the use of RNAi) and reduce the effects of pathogens such as *C*Las.

## Data availability

The datasets supporting this article are available in the *GigaScience* GigaDB repository .

The gene models will be part of an updated official gene set (OGS) for D. citri that will be submitted to NCBI. The OGS (v3) will also be publicly available for download, BLAST analysis and expression profiling on Citrusgreening.org and the Citrus Greening Expression Network [36]. The D. citri genome assembly (v3), OGS (v3) and transcriptomes are accessible on the Citrusgreening.org portal [73]. Accession numbers for genes used in multiple alignments or phylogenetic trees are provided in Table 1.

## Declarations

### List of Abbreviations

ADP: adenosine diphosphate; *Am*: *Apis mellifera*; *Ap*: *Acyrthosiphon pisum*; ATP: adenosine triphosphate; BLASTp: protein BLAST; CGEN: Citrus Greening Expression Network; *C*Las: *Candidatus* Liberibacter asiaticus; *Cmac*: *C. macrophylla; Cmed*: *C. medica*; *Cret*: *C. reticulata*; *Csin*: *C. sinensis*; *Dc*: *Diaphorina citri*; DHAP: dihydroxyacetone phosphate; *Dm*: *Drosophila melanogaster*; *FBPase*: *fructose-1,6- bisphosphatase*; GAP: glyceraldehyde-3-phosphate; *GAPDH*: *glyceraldehyde 3- phosphate dehydrogenase*; *G6Pase/G6P*: *glucose-6-phosphatase*; *Hh*: *Halyomorpha halys*; *HK*: *hexokinase*; Iso-seq: Isoform sequencing; MCOT: Maker, Cufflinks, Oasis, Trinity; NADH: nicotinamide adenine dinucleotide (reduced form); NAD+: nicotinamide adenine dinucleotide (oxidized form); NCBI: National Center for Biotechnology Information; *Nl*: *Nilaparvata lugens*; *PC*: *pyruvate carboxylase*; *PEPCK*: *phosphoenolpyruvate carboxykinase*; *PFK*: *phosphofructokinase*; *PGAM*: *phosphoglycerate mutase*; *PGI*: *phosphoglucose isomerase*; *PGK*: *phosphoglycerate kinase*; *PYK*: *pyruvate kinase*; RNAi: RNA interference; RNA-seq: RNA sequencing; *Tc*: *Tribolium castaneum*; *TPI*: *triosephosphate isomerase*; TPM: transcripts per million; *TPP*: *trehalose-6-phosphate phosphatase*; *TPS*: *trehalose-6-phosphate synthase*; TRE: trehalose; TZP: triazophos; T6P: trehalose-6-phosphate.

### Ethical Approval

Not applicable.

### Consent for publication

Not applicable.

### Authors’ Contributions

WBH, SJB, TD and LAM conceptualized the study; TD, SS, TDS and SJB supervised the study; SJB, TD, SS and LAM contributed to project administration; BT, AM, KK, CV, CM, DH and SA conducted investigation; PH, MF-G, NP and SS contributed to software development; PH, MF-G, SS, TDS and JB developed methodology; SJB, TD, WBH and LAM acquired funding; BT, DLH, MRJ, AM and KK prepared and wrote the original draft; TD, SJB, SS, NP, TDS, WBH and JB reviewed and edited the draft.

## Acknowledgements

We would like to thank Helen Wiersma-Koch (Indian River State College) and Thomson Paris (USDA-ARS-Horticultural Research Laboratories) for assistance.

## Appendix

**Table 1:**
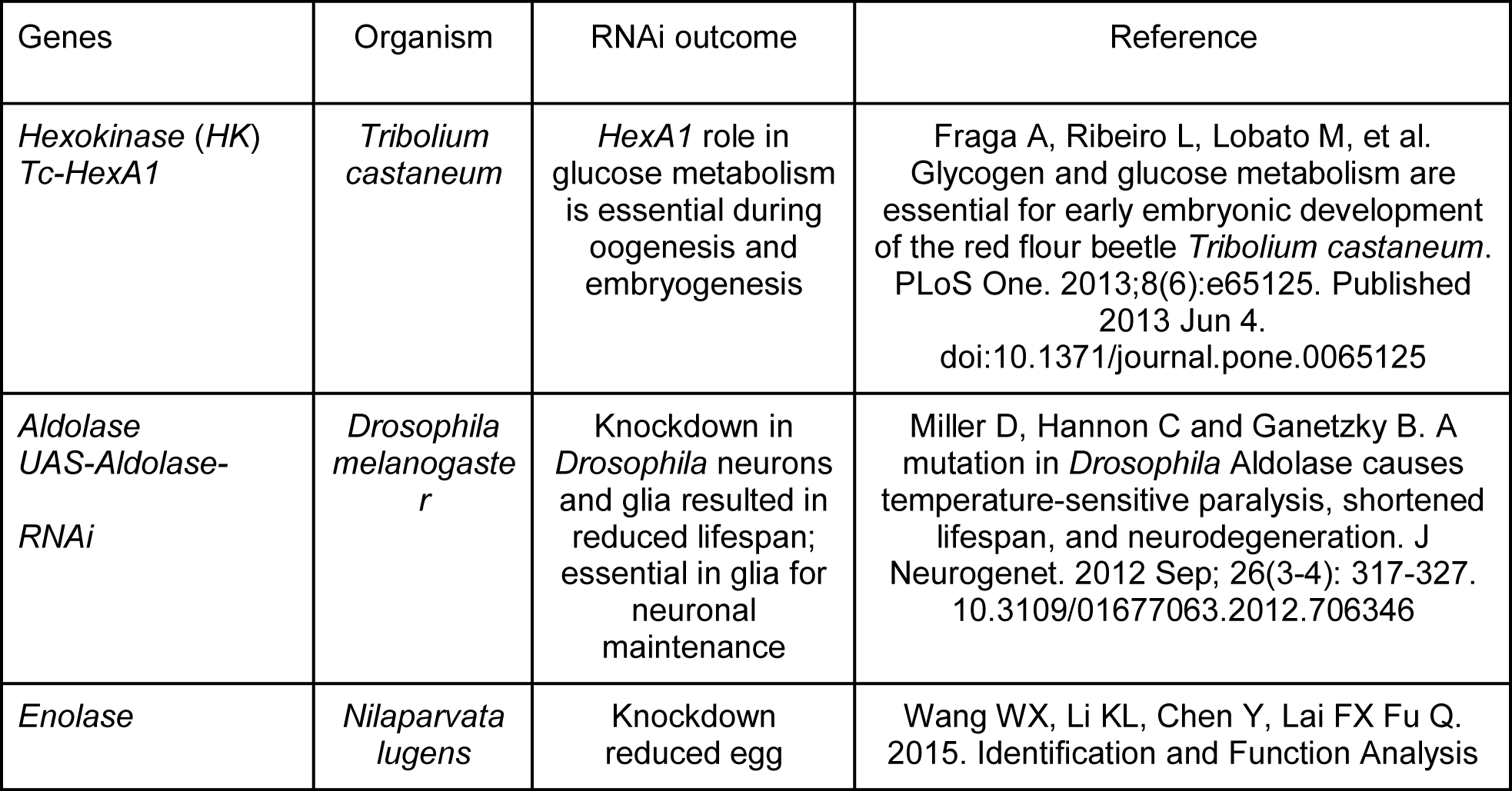
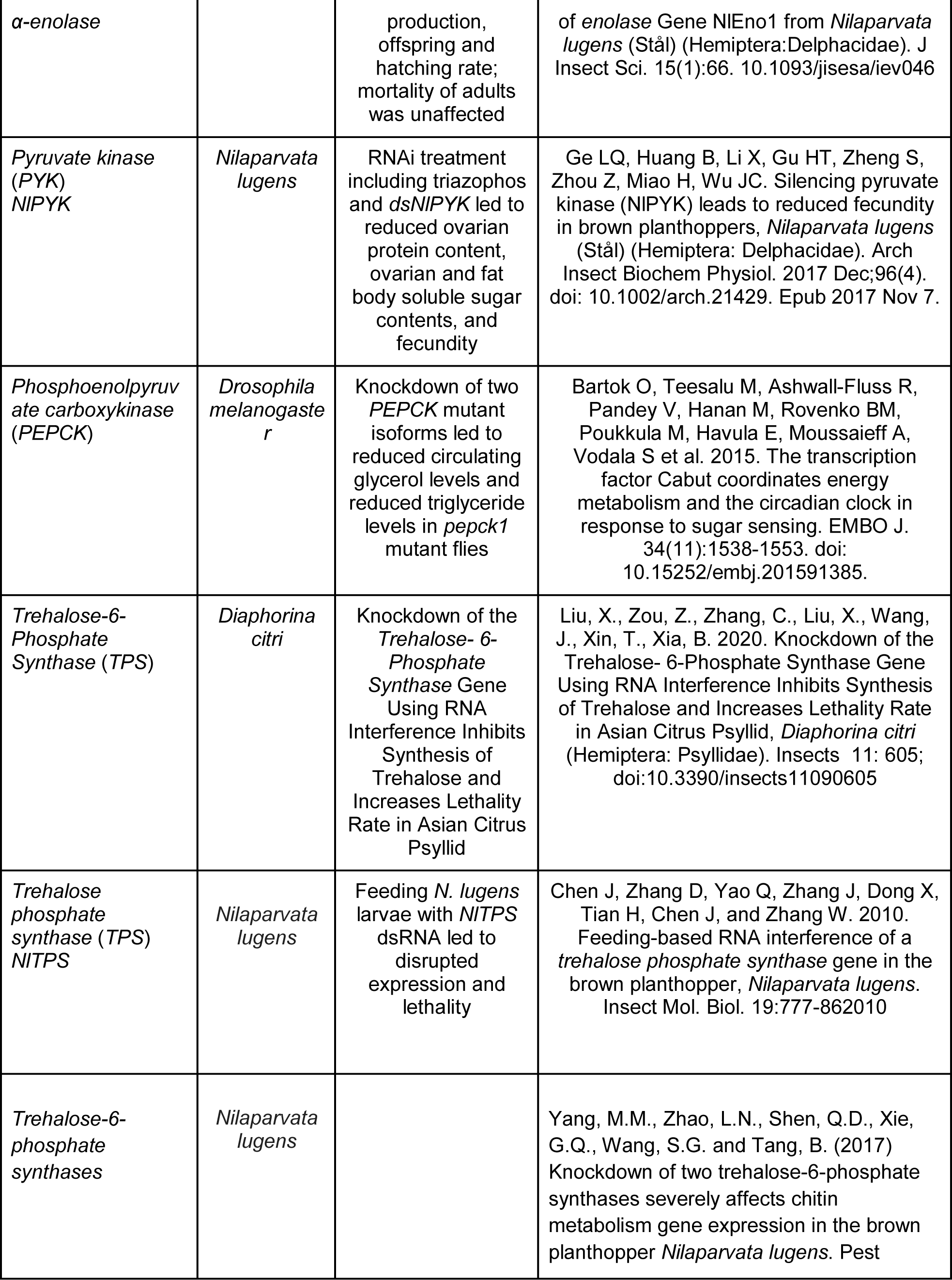
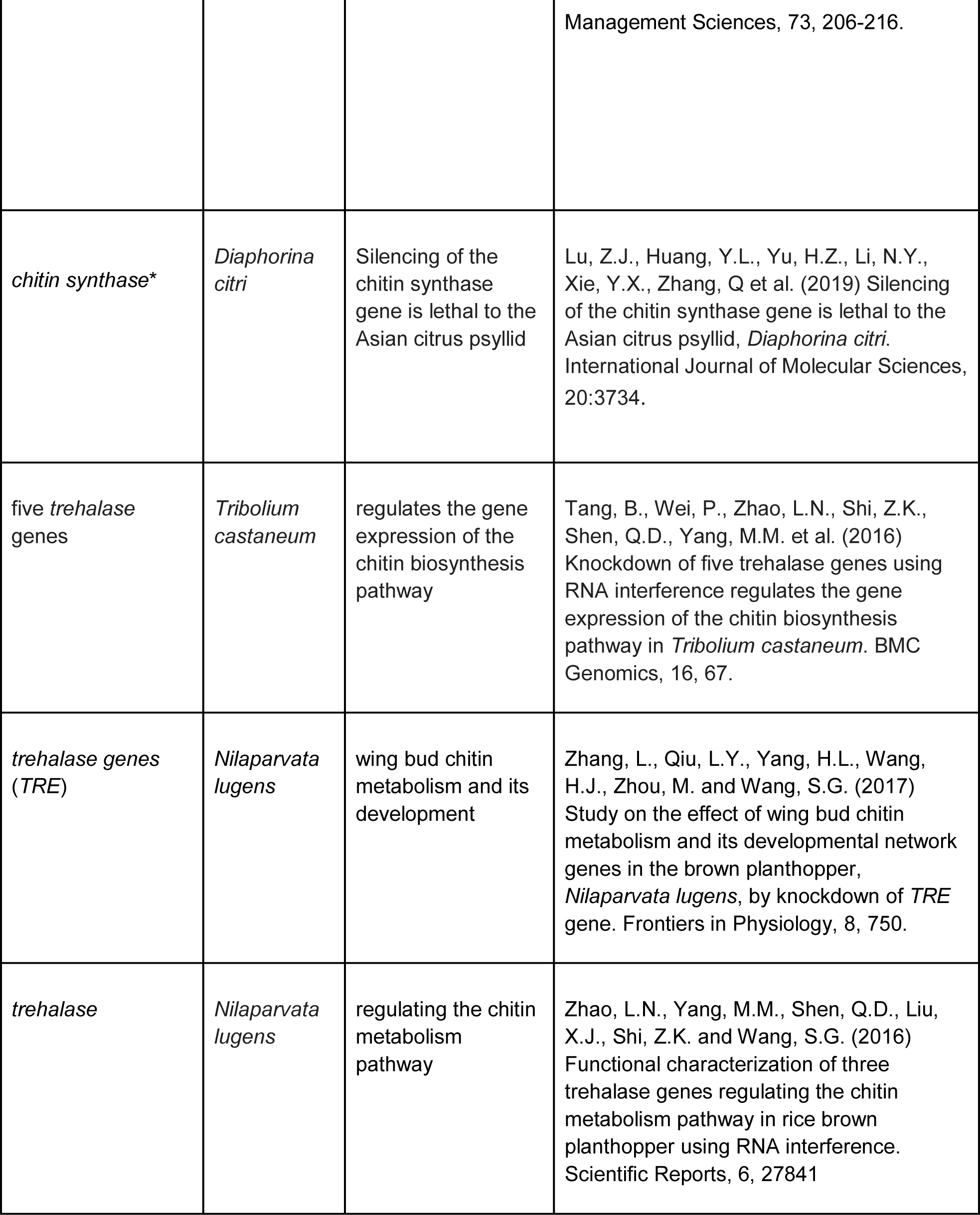
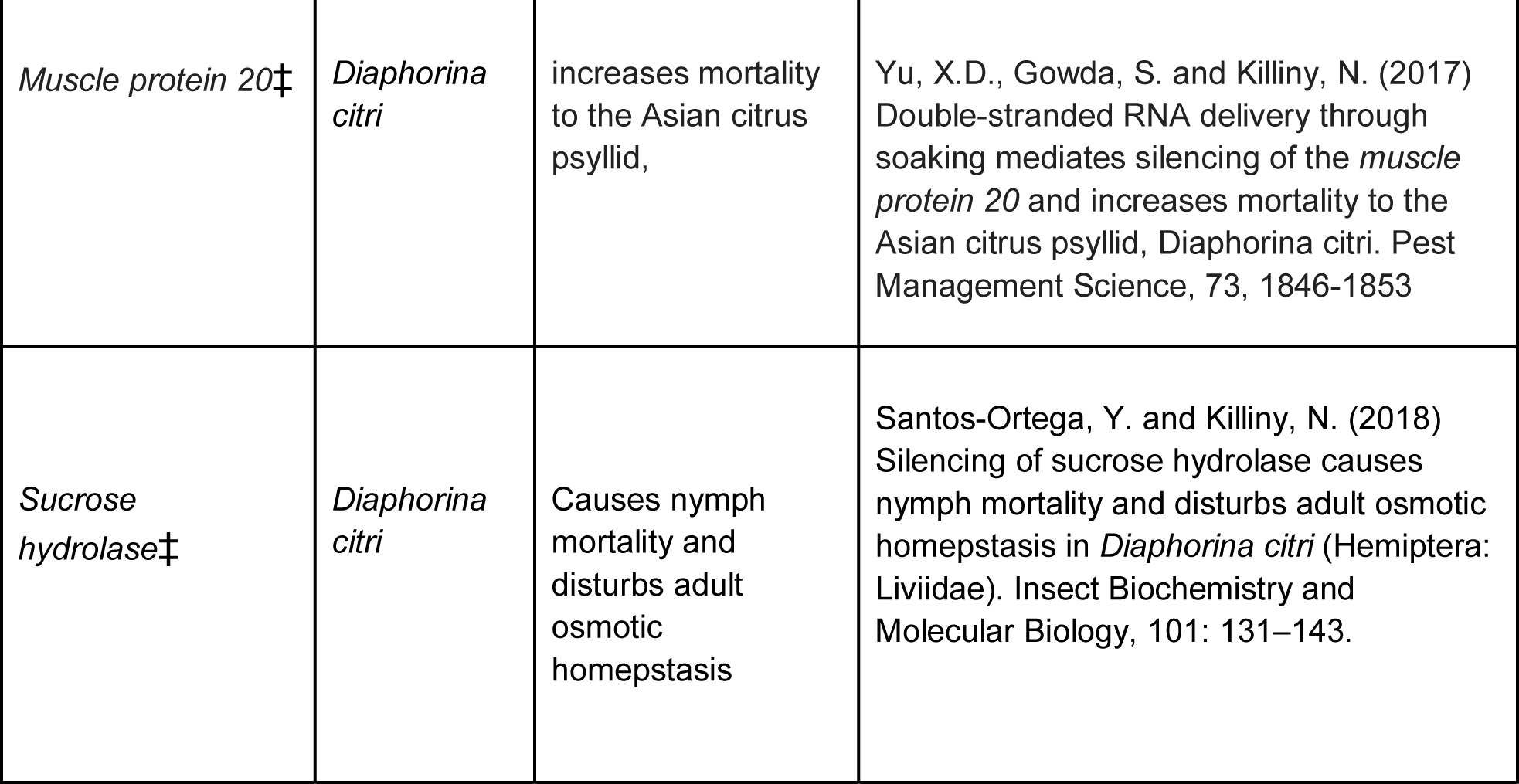
Carbohydrate Metabolism RNAi Gene Targets List of annotated genes in glycolysis (*HK*, *Aldolase*, *Enolase*, *PYK*), gluconeogenesis (*PEPCK*), and trehaloneogenesis (*TPS* and *TREH*), with their corresponding RNAi studies and references. ‡ indicates that additional genes were added, but not annotated in *D. citri*, such as *muscle protein 20* and *sucrose hydrolase*. * indicates that the *chitin synthase* gene in the chitin synthesis pathway was also annotated in *D. citri* [74].

**Table 2:**
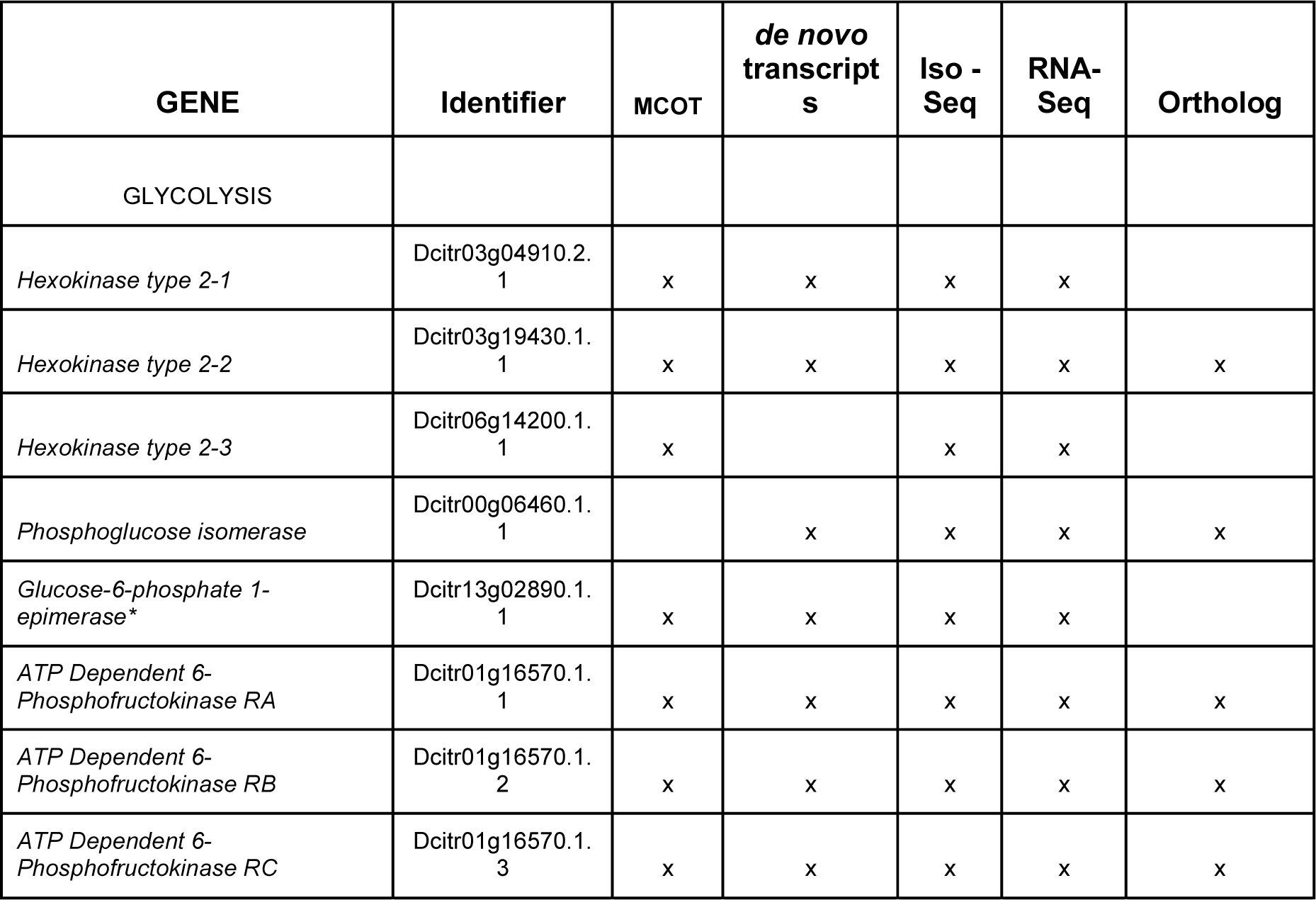
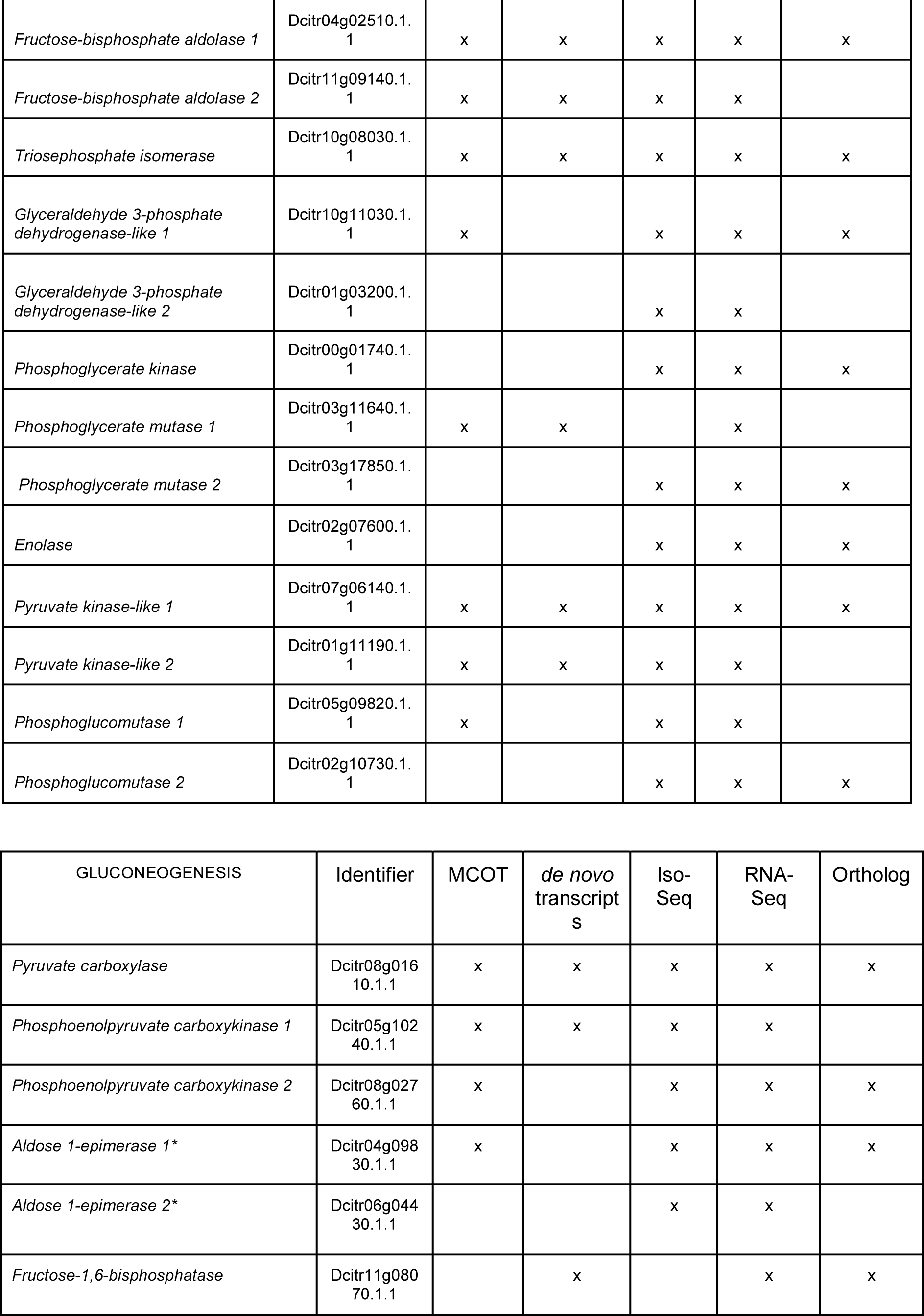
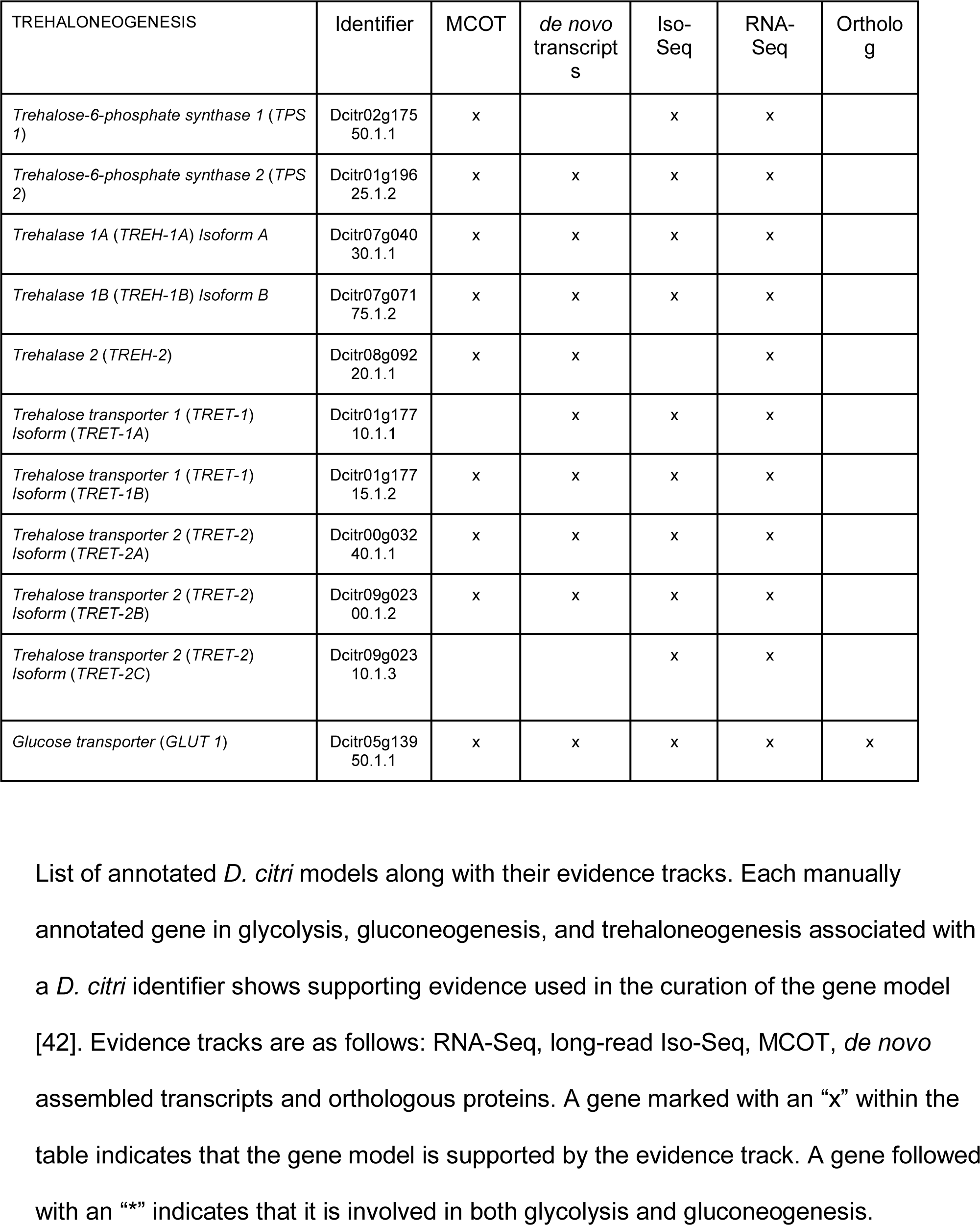
Evidence Table

**Table 3:**
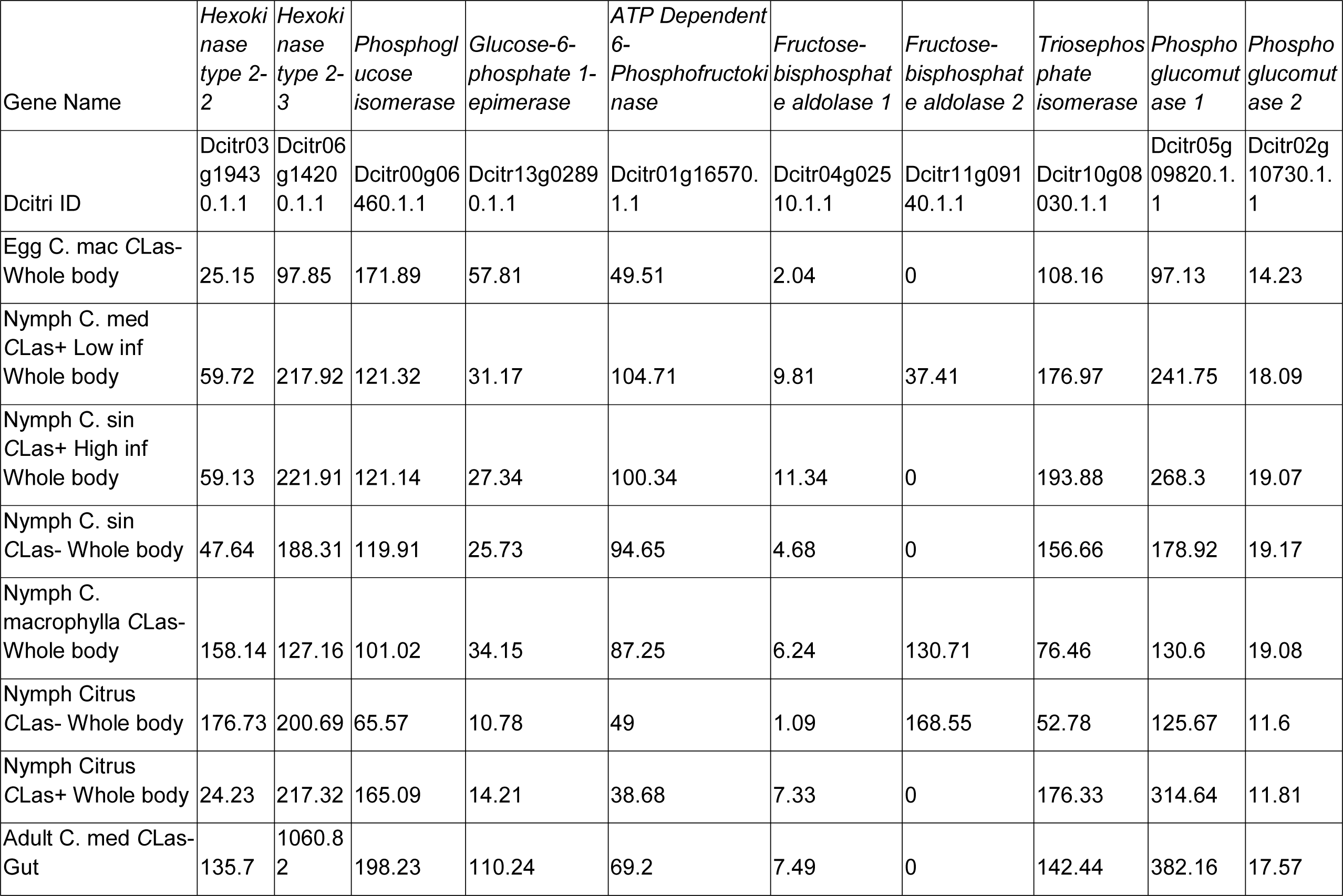
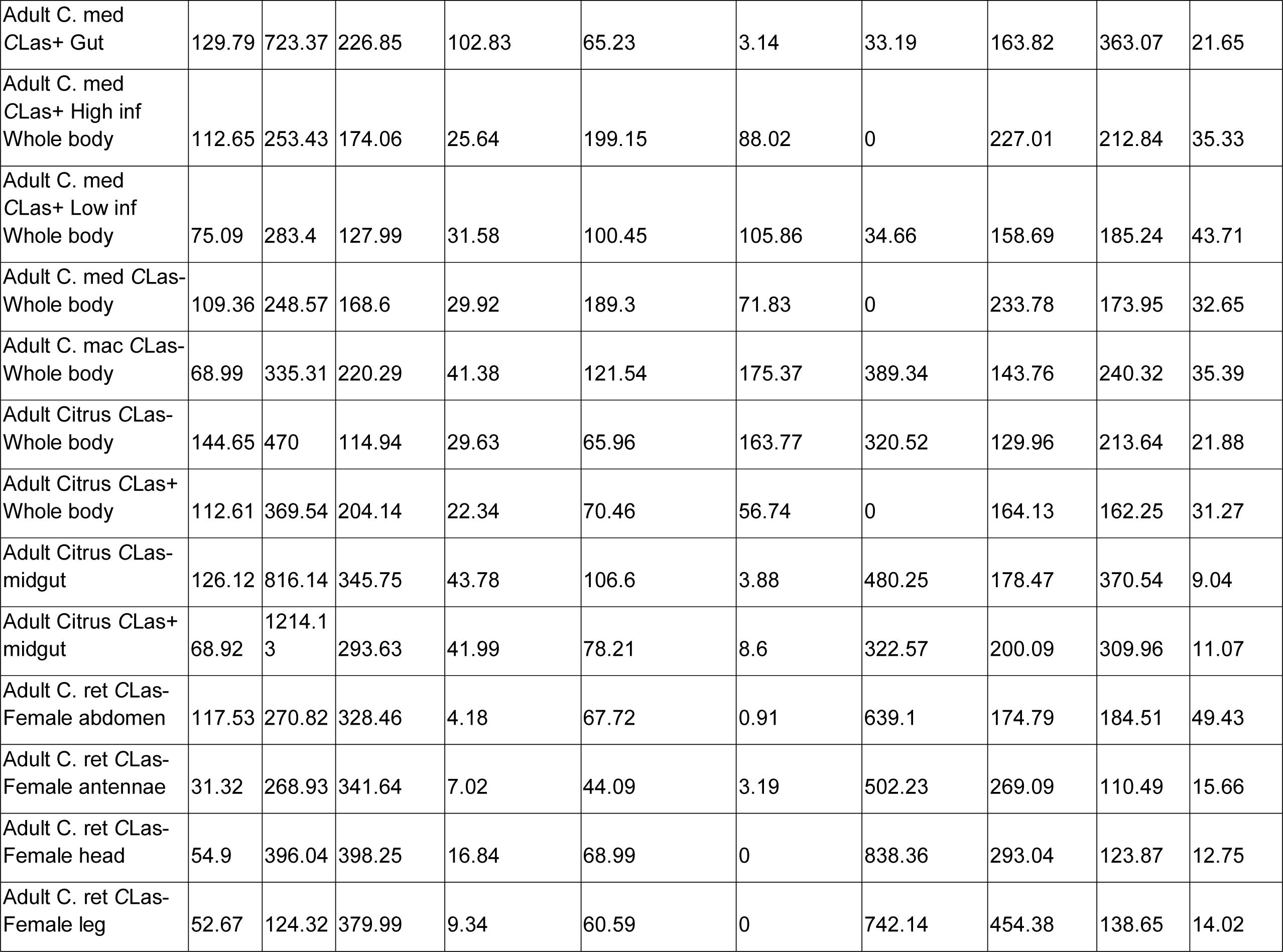
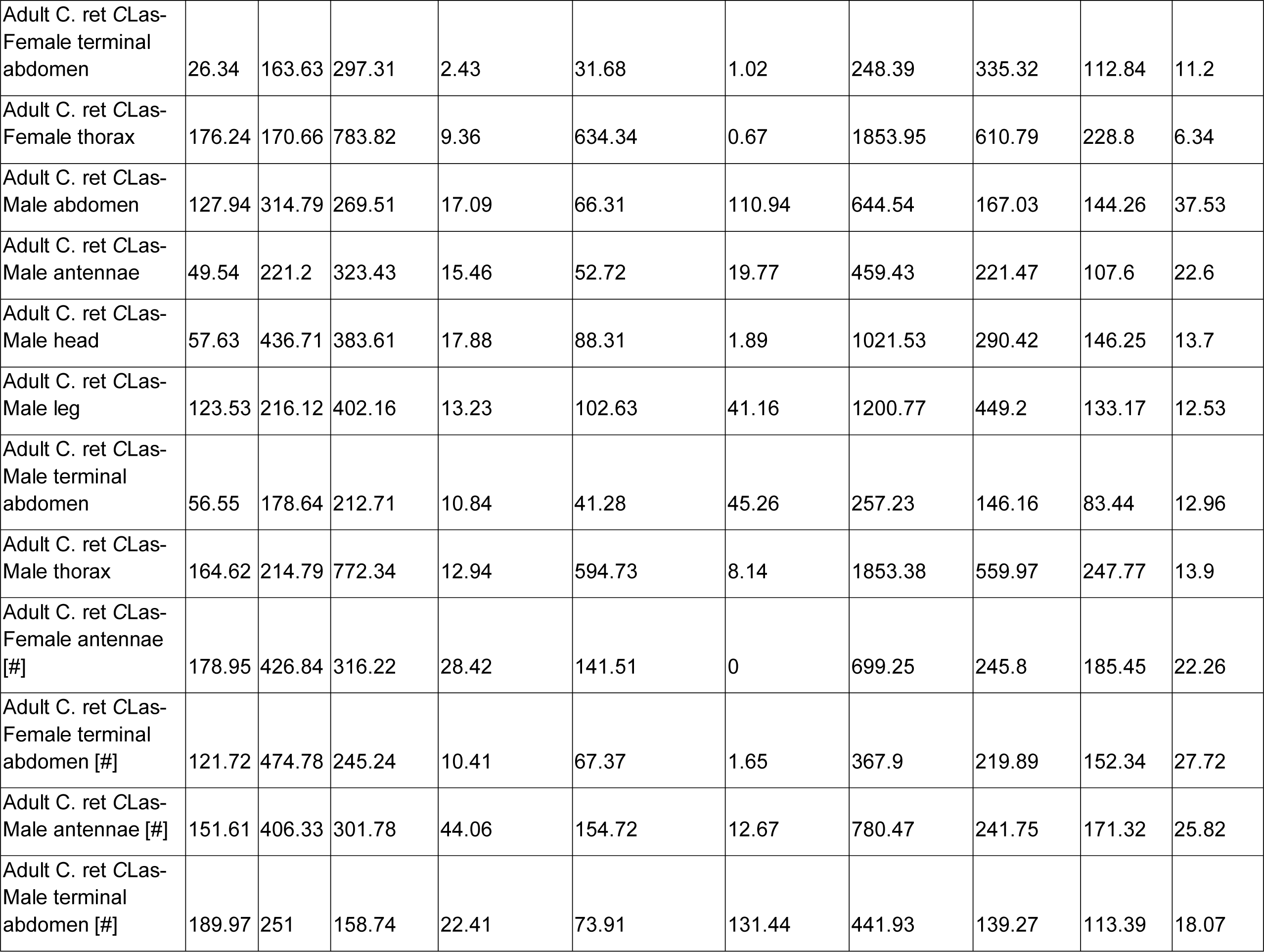
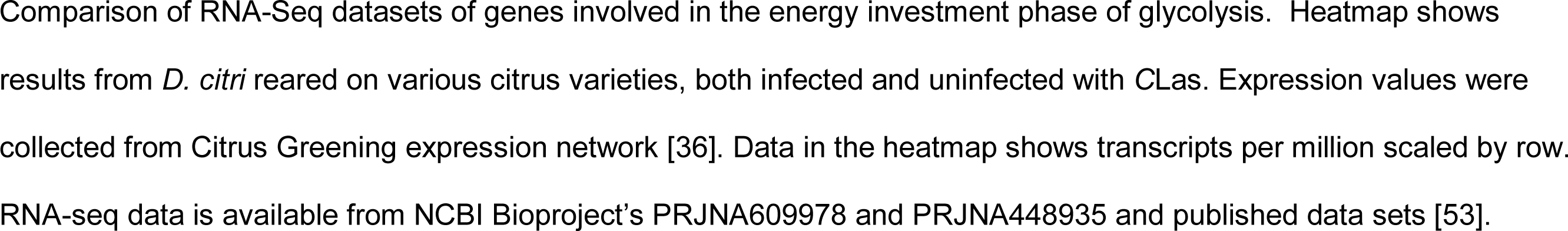
Heat map values in Transcripts per million (TPM); glycolysis energy investment phase

**Table 4:**
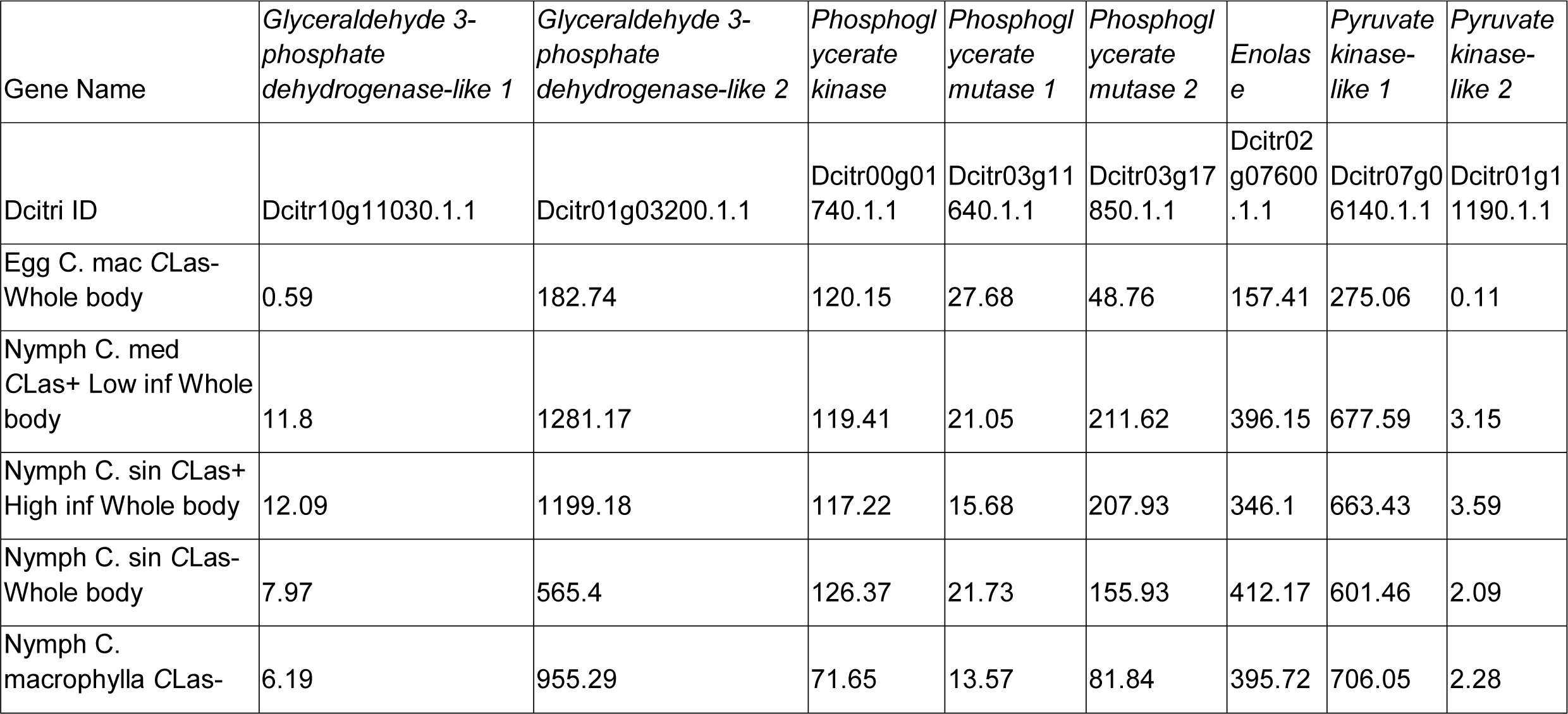
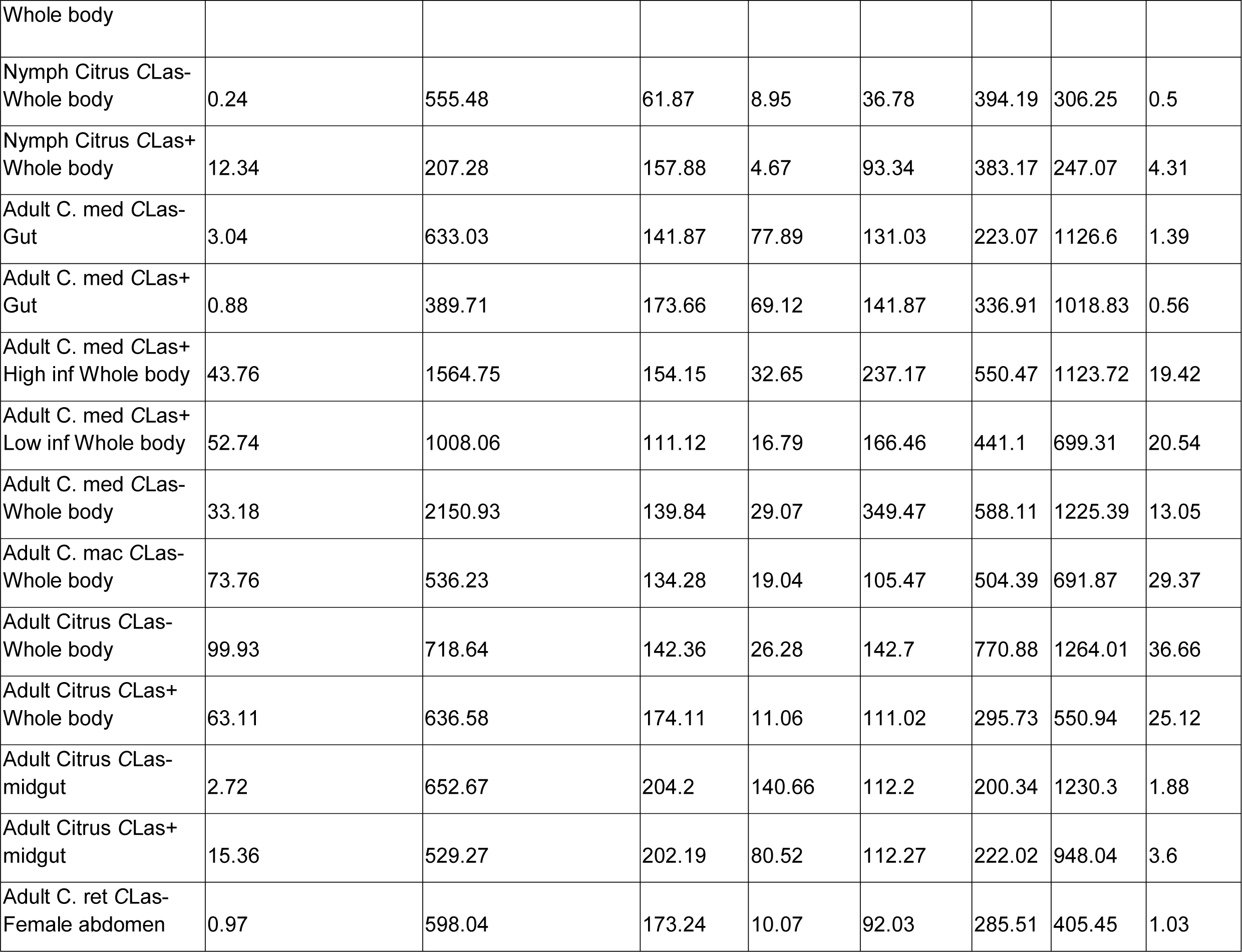
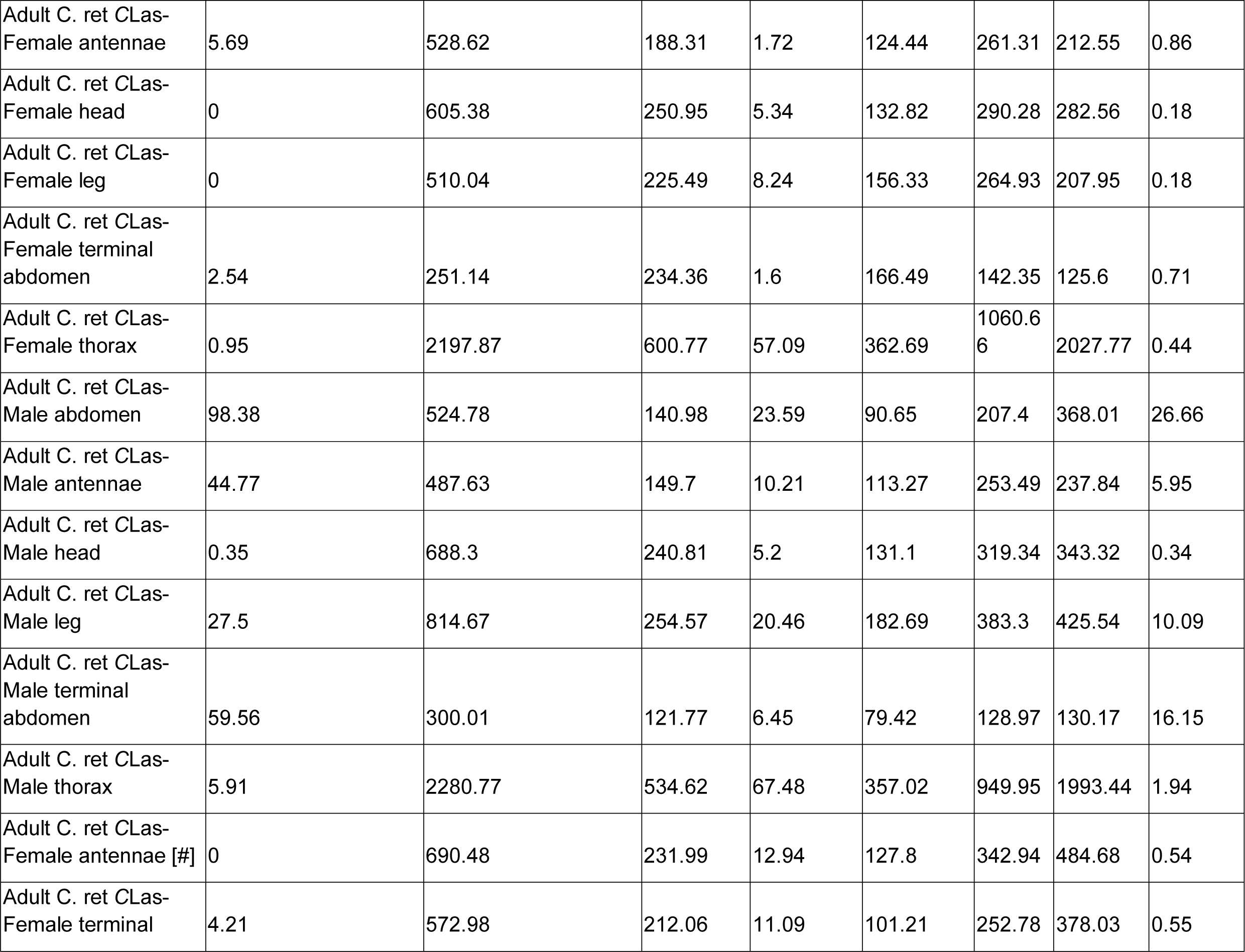
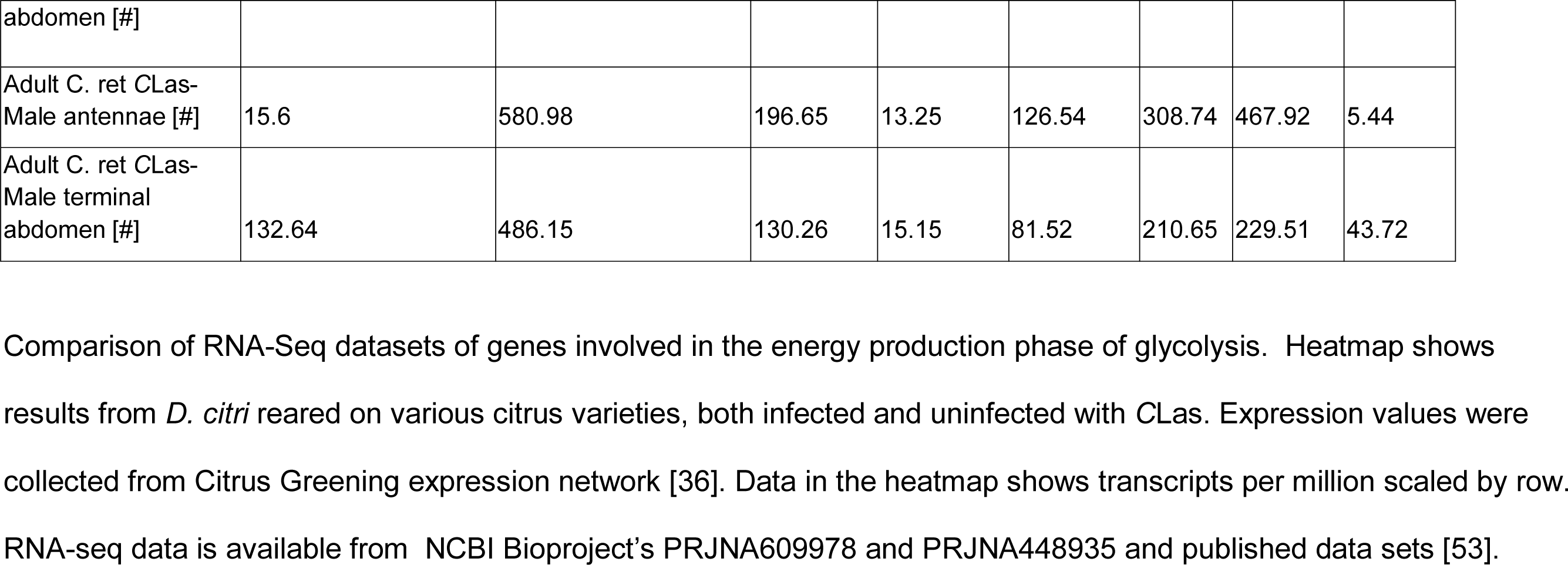
Heat map values in Transcripts per million (TPM); glycolysis energy production phase

**Table 5:**
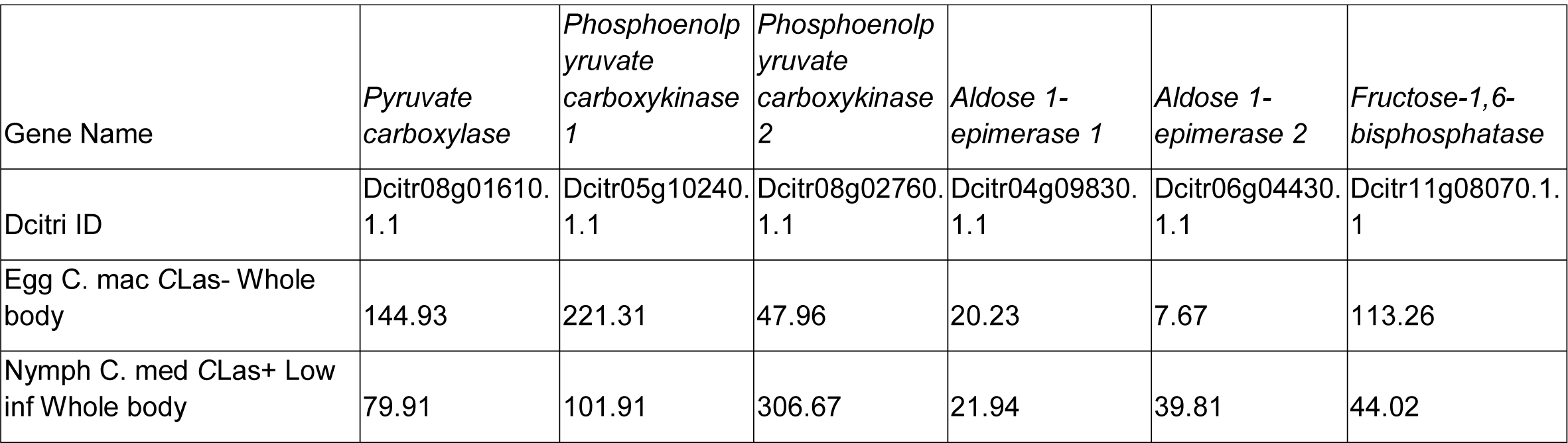
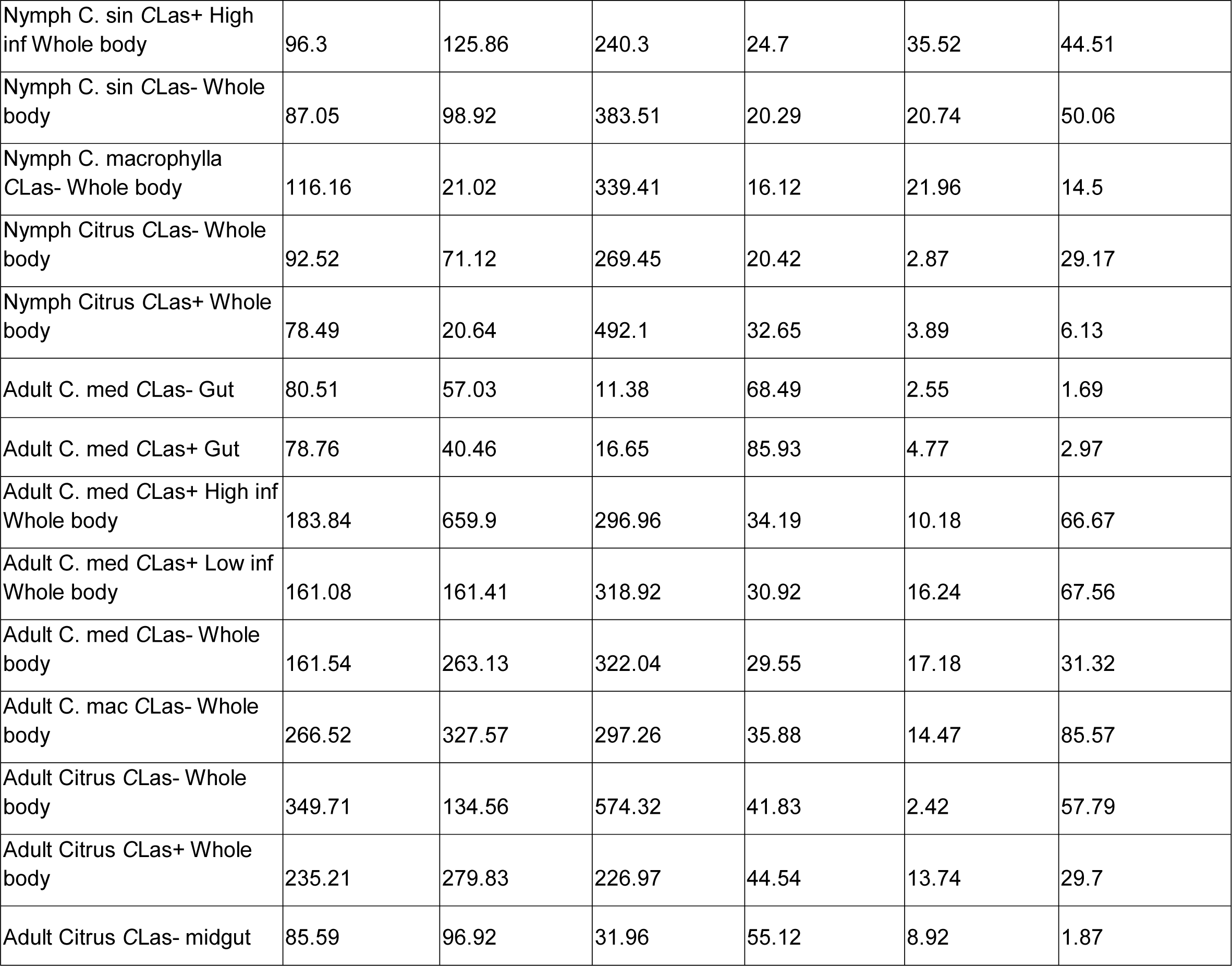
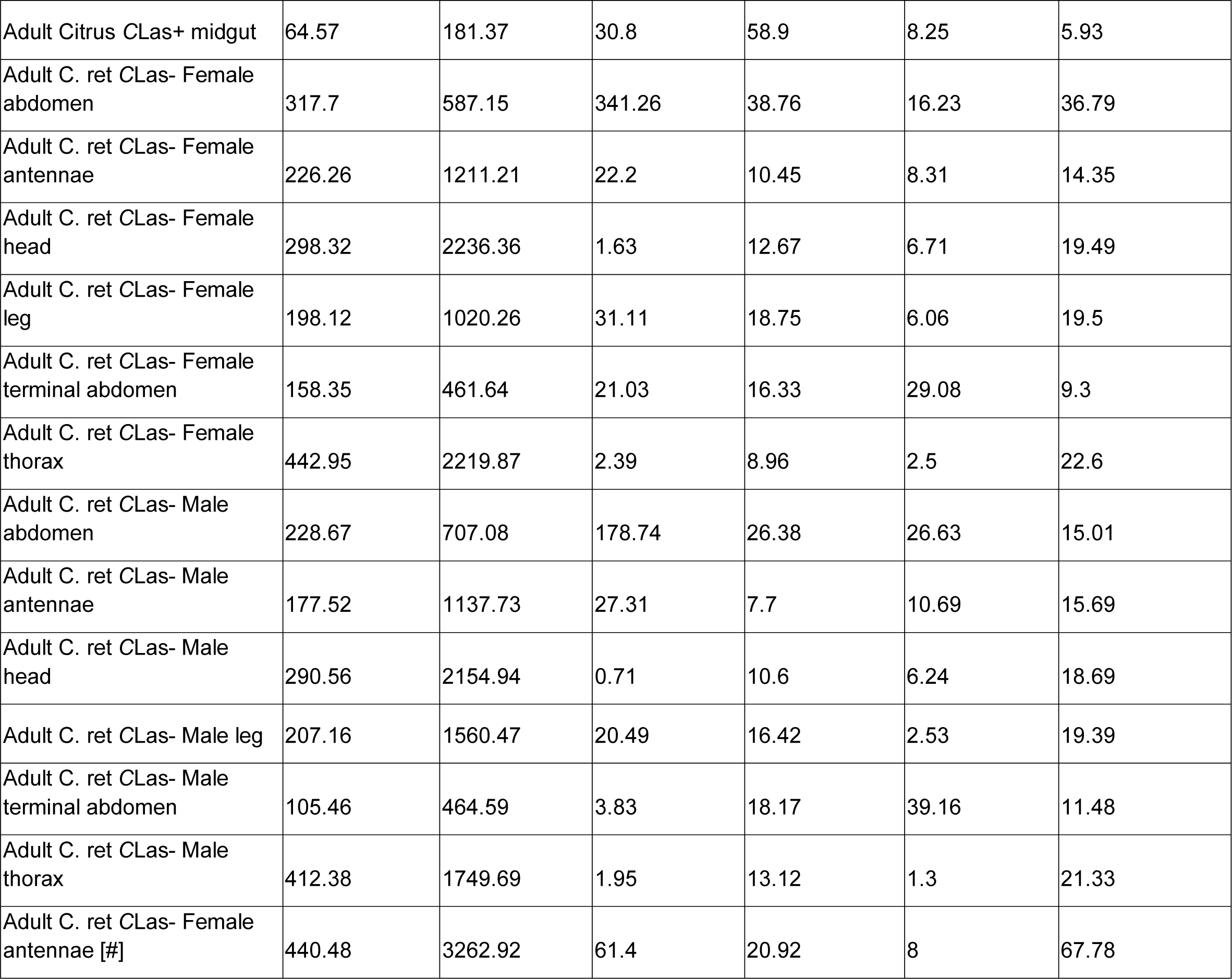
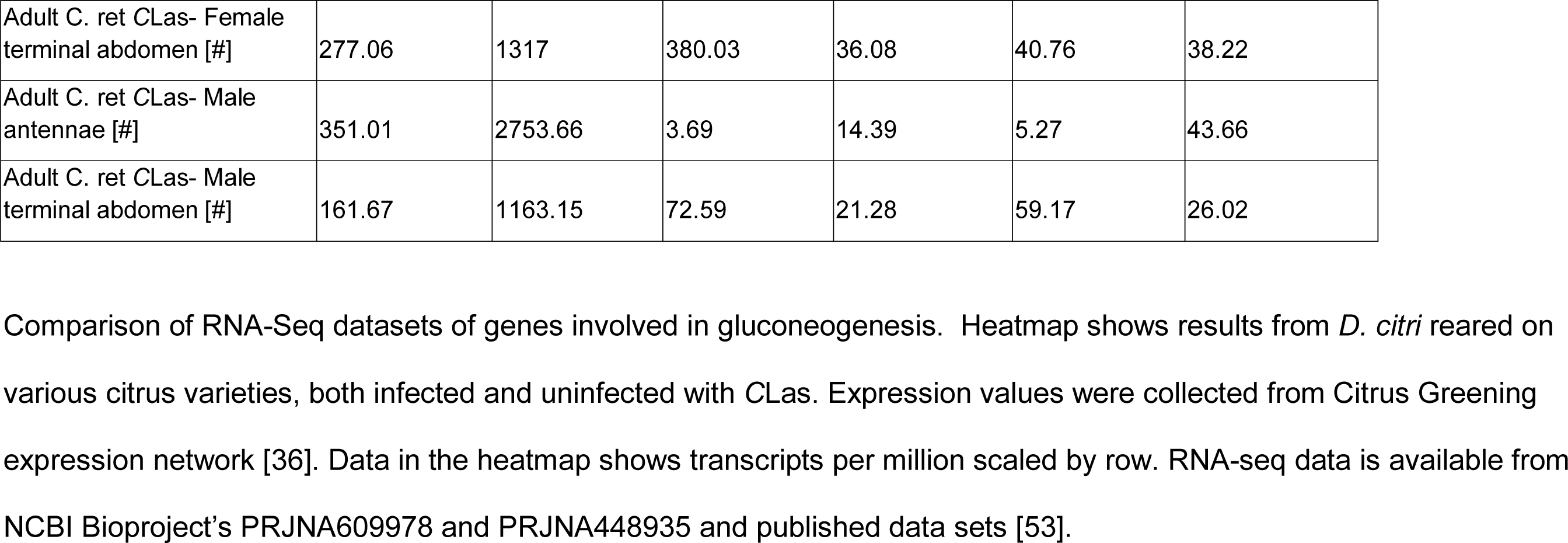
Heat map values in Transcripts per million (TPM); Gluconeogenesis

**Table 6:**
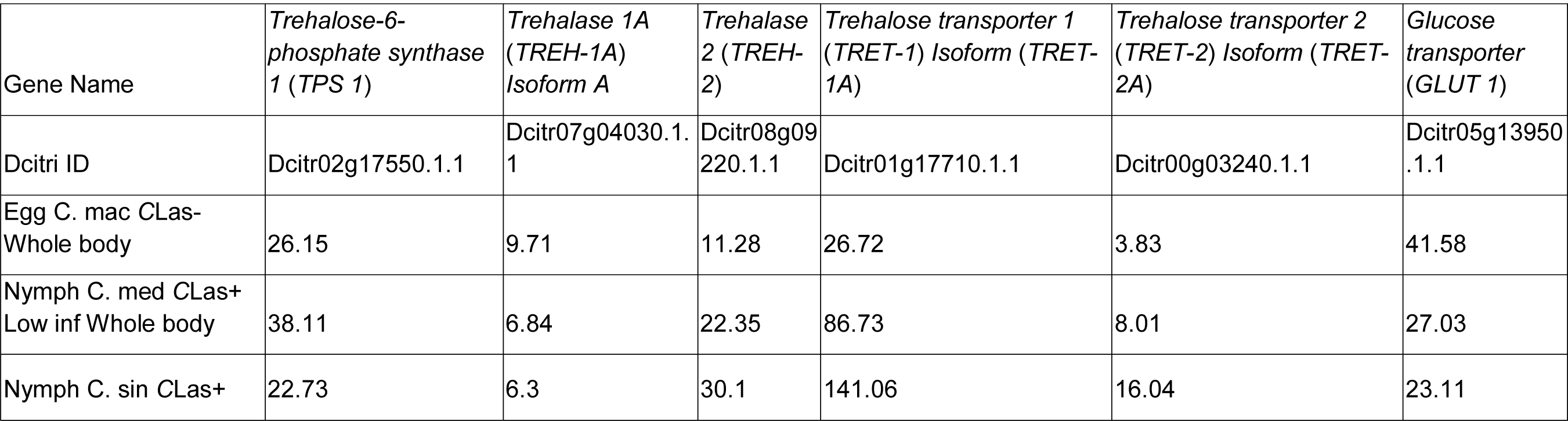
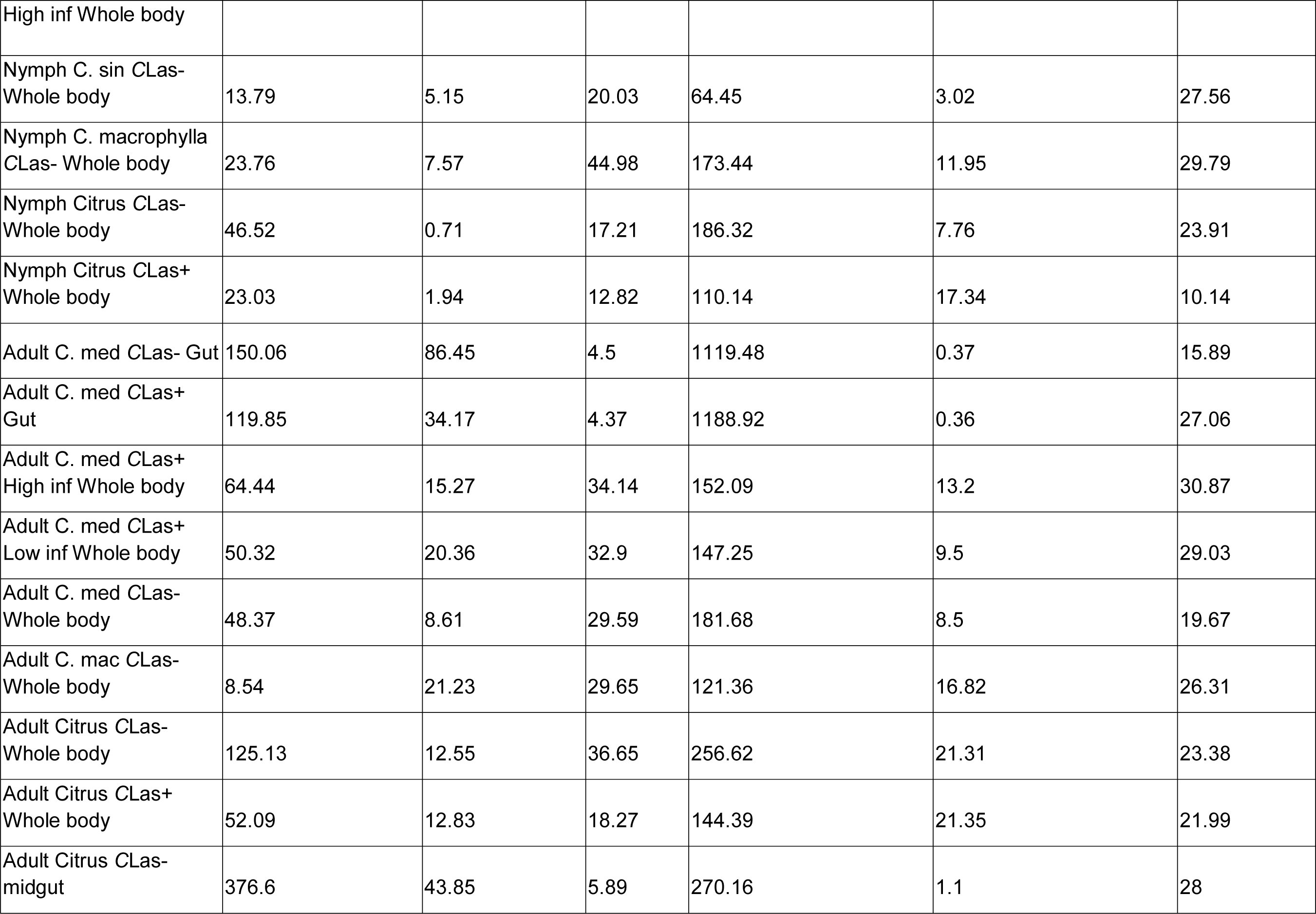
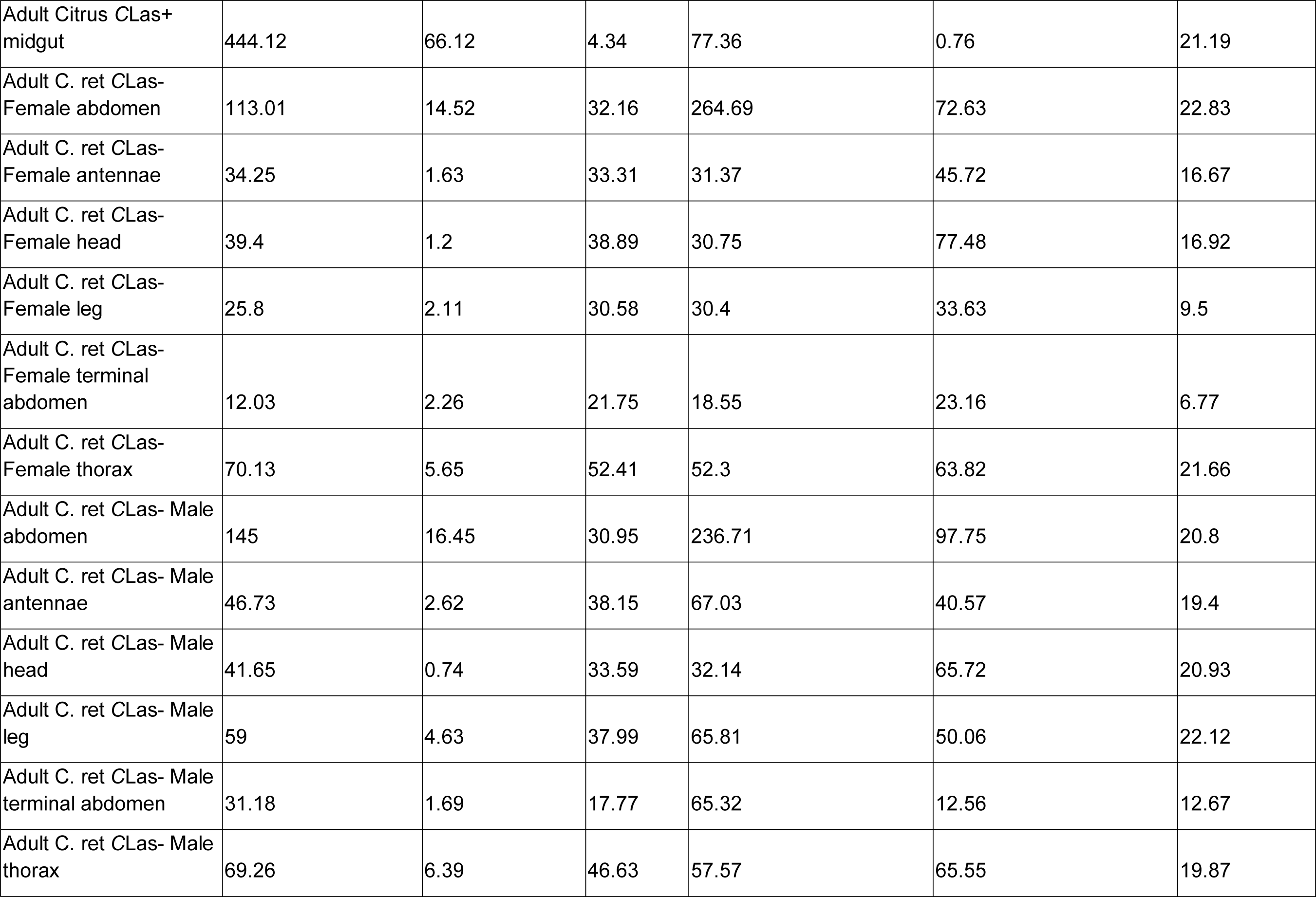
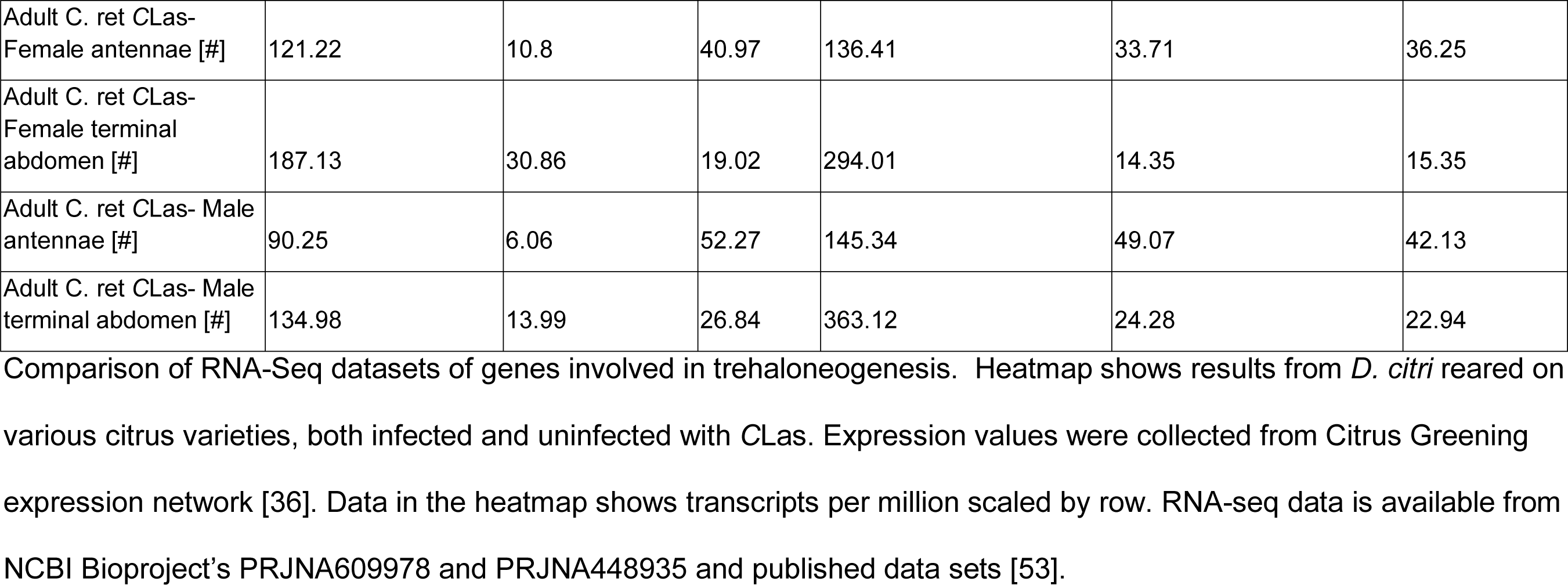
Heat map values in Transcripts per million (TPM); Trehaloneogenesis

